# Synaptic-dependent developmental dysconnectivity in 22q11.2 deletion syndrome

**DOI:** 10.1101/2024.03.29.587339

**Authors:** F.G. Alvino, S. Gini, A. Minetti, M. Pagani, D. Sastre-Yagüe, N. Barsotti, E. De Guzman, C. Schleifer, A. Stuefer, L. Kushan, C. Montani, A. Galbusera, F. Papaleo, M. V. Lombardo, M. Pasqualetti, C.E. Bearden, A. Gozzi

## Abstract

Chromosome 22q11.2 deletion is among the strongest known genetic risk factors for neuropsychiatric disorders, including autism and schizophrenia. Brain imaging studies have reported disrupted large-scale functional connectivity in people with 22q11 deletion syndrome (22q11DS). However, the significance and biological determinants of these functional alterations remain unclear. Here, we use a cross-species design to investigate the developmental trajectory and neural underpinnings of brain dysconnectivity in 22q11DS. We find that LgDel mice, an established mouse model of 22q11DS, exhibit age-specific patterns of functional MRI (fMRI) dysconnectivity, with widespread fMRI hyper-connectivity in juvenile mice reverting to focal hippocampal hypoconnectivity over puberty. These fMRI connectivity alterations are mirrored by co-occurring developmental alterations in dendritic spine density, and are both transiently normalized by developmental GSK3β inhibition, suggesting a synaptic origin for this phenomenon. Notably, analogous hyper-to hypoconnectivity reconfiguration occurs also in human 22q11DS, where it affects hippocampal and cortical regions spatially enriched for synaptic genes that interact with GSK3β, and autism-relevant transcripts. Functional dysconnectivity in somatomotor components of this network is predictive of age-dependent social alterations in 22q11.2 deletion carriers. Taken together, these findings suggest that synaptic-related mechanisms underlie developmentally mediated functional dysconnectivity in 22q11DS.

## Introduction

Complex developmental and neuropsychiatric disorders such as autism and schizophrenia have been consistently associated with dysfunctional interareal communication and atypical functional connectivity ^1, 2^. These observations suggest that brain functional dysconnectivity (i.e., dysfunctional coupling between brain regions) is a systems-level dimension that is key to the pathophysiology of these disorders ^3, 4, 5^. Computational and biophysical models support this view, showing that neuropsychiatric disorders can effectively be conceptualized as (functional) disconnection syndromes ^6, 7^.

Despite the high prevalence of atypical functional connectivity in complex neuropsychiatric disorders, the specific underpinnings of these alterations remain poorly understood. Large-scale functional dysconnectivity may in principle reflect a combination of various multiscale contributing factors, ranging from anatomical miswiring ^8, 9^, to functional and anatomical alterations in neural and synaptic coupling ^10,11^. However, the relative contribution of these components to specific brain disorders, and their relationship to behavior have proven difficult to disambiguate. This problem is compounded by the highly heterogenous etiology and manifestation of brain functional dysconnectivity across patient cohorts ^12, 13^, which together have prevented a reliable decoding of imaging signals into physiologically interpretable events.

Investigations of brain dysconnectivity in genetically-defined neuropsychiatric and developmental disorders may provide valuable mechanistic insight into the underpinnings of these alterations ^8^. A key benefit of this approach is the possibility of relating signatures of functional dysconnectivity to well-characterized etiological factors (and the biological cascade these trigger) in deeply-phenotyped clinical populations. A second important benefit of this approach is the availability, for most of these prevalent syndromes, of animal models that can recapitulate clinically-relevant mutations with high construct validity. Parallel investigations in human cohorts and physiologically accessible animal models can thus be employed to generate testable hypotheses about the contribution (or lack thereof) of specific biological mechanisms to the establishment of brain dysconnectivity.

Chromosome 22q11.2 deletion is among the strongest known genetic risk factors for developmental disorders, including autism and schizophrenia ^14, 15^. The relative high prevalence of 22q11.2 deletion syndrome (22q11DS), estimated to be ca. 1 per 3000 to 1 per 6000 live births in the general population ^16, 17^, has prompted clinical and neuroimaging investigations in increasingly larger aggregated patient cohorts ^18, 19^. Genomic and biological analyses of the ∼46 genes that are typically affected by 22q11.2 deletion, and their modelling in rodents ^20^ have also provided initial clues about some of the putative biological processes underlying psychiatric risk in 22q11DS ^21^. Importantly, recapitulating findings in idiopathic autism and schizophrenia, brain imaging studies have revealed atypical patterns of functional connectivity in individuals with 22q11DS ^22, 23, 24, 25^. However, the neural determinants and significance of the observed functional dysconnectivity remain undetermined. Moreover, the precise developmental trajectory of functional alterations in 22q11DS remains poorly understood. Because 22q11DS is associated with both early and late-onset neurodevelopmental disorders ^14^, gaining a better understanding of the developmental evolution of functional dysconnectivity in 22q11.2 deletion carriers may shed light on the relevance of this phenotype to 22q11DS pathology and specific symptom domains.

Recent progress in mouse brain imaging ^26^ offers the opportunity to generate and test mechanistic hypotheses about the underpinnings of brain dysconnectivity with great mechanistic precision ^11^. Leveraging cross-species neuroimaging data, here we carried out a set of parallel studies in a mouse model of 22q11DS ^27^ and human 22q11.2 deletion carriers aimed to (a) track the developmental trajectory of functional dysconnectivity in this syndrome and (b) probe the biological bases of these alterations. We find that 22q11DS dysconnectivity dramatically reconfigures over puberty, where it mirrors co-occurring developmental alterations in dendritic spine density. We also show that this functional dysconnectivity is predictive of socio-behavioral alterations in 22q11.2 deletion carriers. These findings suggest that synaptic-related mechanisms underlie developmentally-mediated functional dysconnectivity in 22q11DS.

## Results

### Developmental reconfiguration of fMRI dysconnectivity in the LgDel model of 22q11DS

To probe the developmental trajectory of functional dysconnectivity in 22q11DS, we conducted longitudinal resting-state fMRI connectivity mapping in LgDel mice, an established murine model of 22q11DS ^20^. By investigating fMRI connectivity in a rodent model, we sought to tightly control for environmental and genetic variability to a degree not feasible in human clinical research. Because 22q11DS is associated with both pre-pubertal and post-pubertal neurodevelopmental disorders, and given the profound developmental remodeling occurring during this critical time-window, we carried out longitudinal fMRI connectivity mapping in LgDel mice before puberty (i.e. juvenile stage ^28^) and in young adulthood (i.e. post-pubertal stage, postnatal day P35 ± 2 and P111 ± 7, respectively, Figure 1a). To identify regional hubs or hotspots of fMRI dysconnectivity, we computed and compared at the voxel level maps of fMRI global connectivity ^29^, i.e., a measure of mean connectivity of each voxel with the rest of the brain.

**Figure 1.**
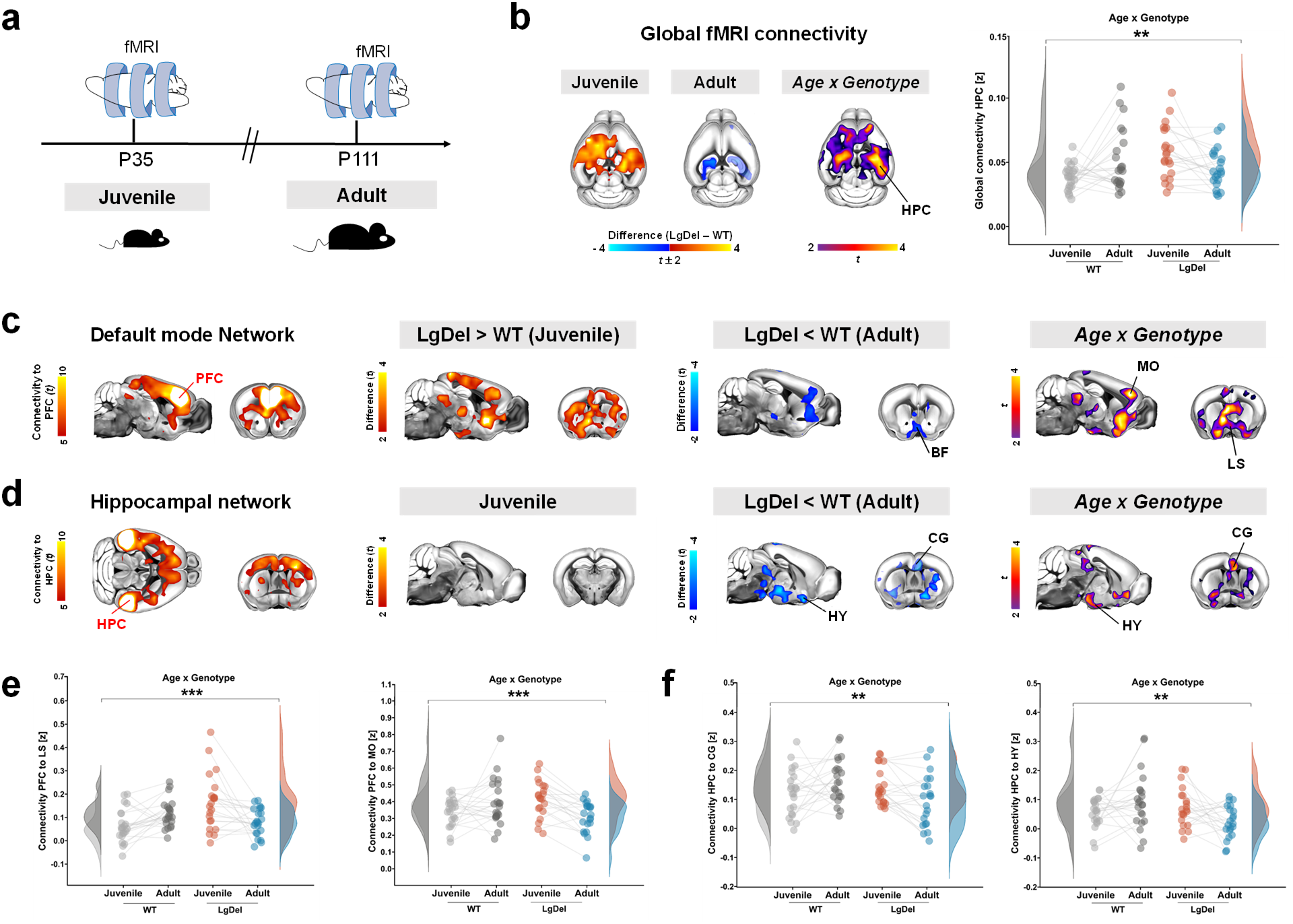
Developmental fMRI dysconnectivity in LgDel mice. (a) Experimental timeline of fMRI mapping in WT and LgDel mice. (b) Whole-brain voxelwise mapping of global fMRI connectivity revealed widespread hyperconnectivity in LgDel juvenile mice, reverting to focal hippocampal hypoconnectivity in the same animals after puberty. Maps are thresholded at |t| > 2.0, followed by FWER correction at p<0.05. Semi-transparent right hippocampal blob indicates a cluster not surviving FWER correction. Corresponding age x genotype maps and quantifications of fMRI global connectivity in dorsal hippocampal areas are reported for reference. (c) and (d) illustrate seed-based mapping of the DMN (c) and Hippocampus (d), respectively. Maps on the left show extension of reference DMN (c) and hippocampal (d) networks in juvenile WT mice. Corresponding between-group difference maps are reported for each of the probed developmental ages in the center panels. Red indicates increased fMRI connectivity, and blue indicates reduced fMRI connectivity compared to control WT littermates (|t| > 2.0, FWER corrected, p < 0.05). (e, f) Regional quantification of pairwise fMRI connectivity between regions of the DMN (e) and hippocampal networks (f) exhibiting significant age x genotype interaction as per maps in (c) and (d). *p<0.05, **p < 0.01, ***p < 0.001. [BF, basal forebrain, CG, Cingulate cortex, HY, hypothalamus, HPC, hippocampus, LS, lateral septum, MO, motor cortex, PFC, prefrontal cortex].

Notably, this investigation highlighted robust alterations of fMRI connectivity in LgDel mice at both the developmental timepoints probed, with evidence of a developmental reconfiguration of the direction and anatomical location of these atypicalities over puberty (Figure 1). Specifically, voxelwise mapping of global fMRI connectivity revealed a switch from widespread hyperconnectivity in juvenile LgDel mice to more focal hippocampal hypoconnectivity in adulthood (Figure 1b, |*t*| > 2.0, *p* < 0.05, FWER cluster-corrected). Corroborating a developmental reconfiguration of fMRI dysconnectivity in LgDel mice, aggregate statistical modelling revealed significant age x genotype interaction in most of the affected regions (i.e., cortex, hippocampus, Figure 1b, |*t*| > 2.0, *p* < 0.05, FWER cluster-corrected).

Seed-based probing of known mouse connectivity networks ^30^ corroborated these findings. Specifically, we found that in juvenile LgDel mice the default-mode network (DMN) (Fig. S1) was functionally hyperconnected to multiple basal forebrain regions (e.g., lateral septum, nucleus accumbens) (Figure 1c). The same regions, however, exhibited fMRI hypoconnectivity (and a significant age x genotype interaction) in the same mice after puberty (Figure 1c, |*t*| > 2.0, FWER cluster-corrected). Similarly, seed-based probing of hippocampal regions did not reveal major connectivity alterations in juvenile LgDel mice, but highlighted robust hypoconnectivity (and significant age x genotype interactions) with the anterior cingulate (a component of the mouse medial prefrontal cortex ^31^ in adulthood, |*t*| > 2.0, FWER cluster-corrected, Figure 1d). These effects (and their age x genotype interactions) were also apparent in subsequent quantifications of pairwise fMRI connectivity in regions of interest (Figure 1e, f). Collectively, these results reveal a previously unreported developmental reconfiguration of fMRI dysconnectivity over puberty in the LgDel model of 22q11DS.

### Developmental reconfiguration of fMRI dysconnectivity in human 22q11.2 deletion carriers recapitulates findings in 22q11.2 mouse model

Prompted by our mouse findings, we next probed whether an analogous reconfiguration of resting-state fMRI) dysconnectivity would also be identifiable over development in individuals with 22q11DS. To this purpose, we aggregated three independent fMRI imaging datasets in 22q11DS (i.e., UCLA, KCL and SUNY), to obtain a single, harmonized cohort of n = 139 22q11.2 deletion carriers (22q11DS) and n = 117 healthy controls (HC), spanning an age range of 6 to 30 years-old (Table 1 and Supplementary Table 1). To replicate the data analysis strategy we used in the mouse, we split the clinical cohort into two age subgroups (Figure 2a). Age ranges were chosen to broadly recapitulate corresponding developmental stage in rodents. We thereby defined a first age group that we herein refer to as *Childhood* (age range 6-11 years old; 22q11DS n = 21, HC n = 31) and a second age group we term *Post-puberty* (age range 12-30 years old; 22q11DS n = 118, HC n = 86). We chose the age cutoff based on previous work pointing at age 11 being a peak of puberty onset (i.e., increased incidence of Tanner stages II/III ^32^).

**Figure 2.**
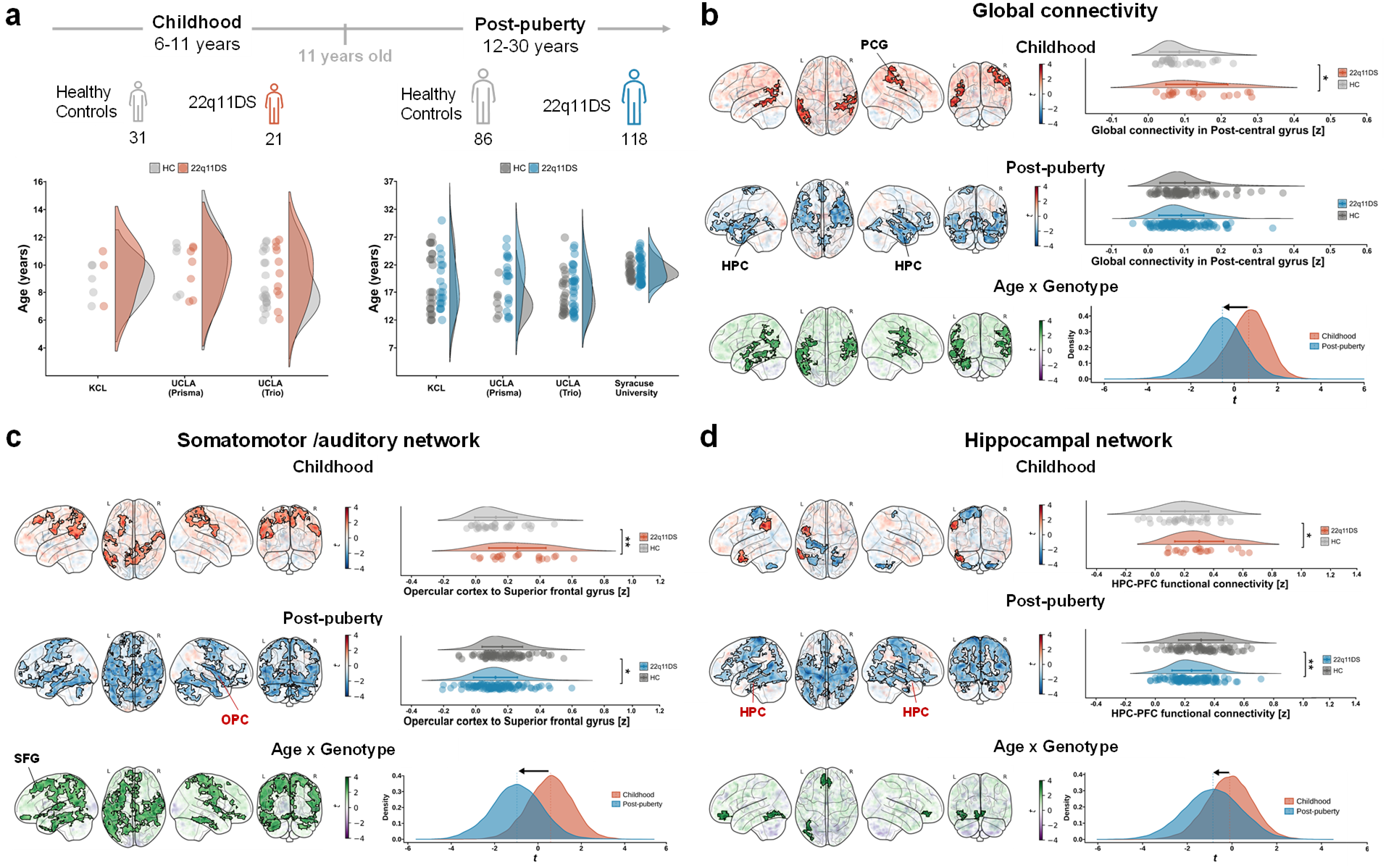
Developmental fMRI dysconnectivity in human 22q11DS. (a) Schematic representation of human cohort divided into the Childhood cohort (HC n=31; 22q11DS n=21) and Post-pubertal cohort (HC n= 86; 22q11DS n=118). Distribution of age for each diagnosis (HC and 22q11DS) across sites and scanners. (b) Voxel-wise (left panels) mapping of global fMRI connectivity revealed increased functional connectivity in 22q11DS carriers during childhood, and reduced fMRI connectivity in the post-pubertal cohort. Semi-transparent maps in the background represent unthresholded t-values. Clusters surviving FWER correction are outlined in black. Areas exhibiting a significant age x genotype interaction were identified using a linear model. Whole-brain distribution of t values resulting from group differences at each age revealed a robust shift (arrow) from prevalent hyperconnectivity in childhood to prevalent hypoconnectivity after puberty (bottom right of panel). (c) and (d) Seed-based analysis using clusters exhibiting significant age x genotype interaction in our global fMRI connectivity analysis as seeds: OPC (c) and hippocampus (d). Distribution of t values resulting from between-group connectivity differences over development are also reported for reference. Note shift from hyper-to hypoconnectivity (arrow) occurring over puberty. [HPC, Hippocampus, OPC, Opercular cortex]. * p < 0.05, ** p < 0.01, *** p < 0.001.

**Table 1.** Subject demographics.

|  | Children<br>(6-11 years) |  |  | Post-pubertal<br>(12-30 years) |  |  |
| --- | --- | --- | --- | --- | --- | --- |
|  | HC<br>(n=31) | 22q11DS<br>(n=21) | *p* value | HC<br>(n=86) | 22q11DS<br>(n=118) | *p* value |
| Age (years) | 8.85 ± 2 | 9.4 ± 2 | 0.26 | 18.4 ± 4 | 19.4 ± 4 | 0.11 |
| Sex (M/F) | 15/16 | 9/12 | 0.69 | 41/45 | 54/64 | 0.79 |
| Full scale IQ* (mean ± SD) | 117.3± 22.6 | 77.7± 7.2 | <0.0001 | 109.4± 16.4 | 79.9± 13.9 | <0.0001 |
| ASD, n (%) | 0 (0) | 7 (33) | 0.0005 | 0 (0) | 16 (13.5) | 0.0004 |
| ADHD, n (%) | 1 (3) | 10 (48) | 0.0001 | 7 (8) | 36 (30) | 0.0001 |
| Psychotic disorder, n (%) | 0 (0) | 0 (0) | - | 0 (0) | 7 (5.9) | 0.02 |
| Anxiety disorder, n (%) | 1 (3) | 7 (33) | 0.0032 | 10 (12) | 30 (25) | 0.02 |
| Mood disorder, n (%) | 0 (0) | 2 (9.5) | 0.08 | 4(5) | 29 (25) | <0.0001 |
| Current antipsychotic treatment, n (%) | 0 (0) | 1 (3) | 0.22 | 0 (0) | 14 (12) | 0.0009 |
| Current antidepressant/anti-anxiety treatment, n (%) | 0 (0) | 0 (0) | - | 2 (2.3) | 24 (20) | 0.0002 |
| Current psychostimulant treatment, n (%) | 1 (3) | 1 (3) | 0.22 | 3 (3.5) | 13 (11) | 0.05 |

\* IQ data were not available for 1 child HC, 2 children 22q11DS, 9 post-pubertal HC, 2 post-pubertal 22q11DS.

As in our mouse investigations, we first carried out spatially-unbiased global fMRI connectivity mapping in the two groups of participants (i.e., 22q11DS, healthy controls), at both ages. Paralleling our findings in LgDel mice, this analysis revealed a clear reconfiguration from hyper-to hypoconnectivity over puberty in people with 22q11DS (Figure 2b). Specifically, in prepubertal 22q11 deletion carriers we found extended foci of global fMRI hyperconnectivity in postero-temporal and somatomotor regions, as well as in the dorsal attention network. By contrast, post-pubertal 22q11.2 deletion carriers exhibited robust fMRI hypoconnectivity relative to controls in the hippocampus and temporal lobe regions (|*t*|>2.0, FWER cluster-corrected). As seen in LgDel mice, most of the affected regions also exhibited robust age x genotype interactions (|*t*|>2.0, FWER cluster-corrected, Figure 2b), thus supporting the notion of an altered developmental trajectory in 22q11DS.

To probe the brain networks most affected by these alterations, we conducted a targeted seed-based analysis of the regional clusters that exhibited a statistically significant age x genotype interaction in our global connectivity mapping (seed location in Figure S2). Seed-probing of a cluster located in the right opercular cortex (a component of the somatomotor/auditory network^33^) showed that prepubertal individuals with 22q11DS exhibited an aberrant extension of this network extending beyond somatomotor regions to include regions belonging to the DMN (middle frontal gyrus, cingulate gyrus, precuneus) and dorsal attention network (lateral occipital cortex, Figure 2c). After puberty, the same network appeared to be largely hypoconnected in 22q11DS. Notably, analogous findings were obtained when probing fMRI connectivity of an age x genotype interaction cluster located in the postcentral gyrus (Figure S3a). We also conducted seed-based probing of the hippocampus (Figure 2d), to test whether post-pubertal 22q11DS carriers would exhibit hypoconnectivity of this region, as found in mice. This analysis showed that focal hippocampal dysconnectivity in pre-pubertal 22q11.2 deletion carriers evolves into robust and widespread hypoconnectivity after puberty. Mirroring mouse findings, the medial prefrontal cortex was amongst the regions exhibiting a transition from hyperconnectivity to hypoconnectivity (i.e., a significant age x genotype interaction, |t| > 2.0, FWER cluster-corrected corrected) during development (Figure 2d, Figure S3b).

Lastly, we assessed whether considering age as a continuous variable would reveal a trajectory of developmental dysconnectivity broadly consistent with our *a priori* categorical analysis of prepubertal and post-pubertal cohorts. We thus employed a non-linear mixed model (GAMM^34^) to probe the relationship between age and fMRI global connectivity in cortical regions undergoing developmental reconfigurations within the somatomotor/auditory network (i.e., opercular cortex, postcentral gyrus), and in the hippocampus. We found the corresponding trajectories of dysconnectivity to be broadly consistent with our age cutoff. Specifically, hyperconnectivity persisted until age 9.2 and 9.9 years in opercular cortex and postcentral gyrus, respectively. Reversal to hypoconnectivity was observed in opercular regions at age of 13.8 years. In the hippocampus, we found the onset of hypoconnectivity to occur at age 12.8 years (Figure S4). These results support our use of a cutoff developmental point of 11 years of age. More broadly, our investigations document the presence of developmental fMRI dysconnectivity in 22q11DS, involving a transition between hyperconnectivity to hypoconnectivity relative to typically developing controls across puberty.

### Developmental dysconnectivity in LgDel mice is associated with GSK3β-dependent alterations in dendritic spine density

The observed correspondence between human and mouse neuroimaging findings prompted us to use the LgDel model to investigate the neurobiological underpinnings of developmental fMRI dysconnectivity in 22q11DS. Neuroanatomical investigations have provided evidence of synaptic-related pathology and altered dendritic spine density in LgDel mice and other rodent models of 22q11DS, especially in hippocampal areas ^35, 36, 37^. Using *in vivo* manipulations and computational modelling, we recently showed that alterations in dendritic spine density can be putatively mirrored by corresponding changes in global fMRI connectivity, with increased or decreased spine density being paralleled by increased ^10^ and decreased ^38^ global fMRI connectivity, respectively. These observations suggest that synaptic-related mechanisms might contribute to fMRI dysconnectivity in LgDel mice. This hypothesis would be consistent with the profound synaptic remodeling occurring in the mammalian brain during adolescence ^39^.

To test this hypothesis, we first measured dendritic spine density in the PFC and hippocampus of LgDel mice at juvenile and adult stage (Figure 3a, Figure S5). We found that dendritic synaptic density was altered in both the probed regions, albeit with different developmental trajectories. Specifically, spine density in the PFC was robustly increased in juvenile LgDel mice but showed highly comparable values between groups in adulthood (Figure 3a, p < 0.0001 and p > 0.99, respectively, *Sidak’s multiple comparisons test*, two-way ANOVA; Figure S5). In the hippocampus we found a similar increase in dendritic spine density in juvenile LgDel animals. However, a reversed phenotype was observed after puberty, with LgDel mice exhibiting significantly lower spine density than WT littermates (Figure 3a, p < 0.0001 and p = 0.01, respectively, *Sidak’s multiple comparisons test*, two-way ANOVA; Figure S5).

**Figure 3.**
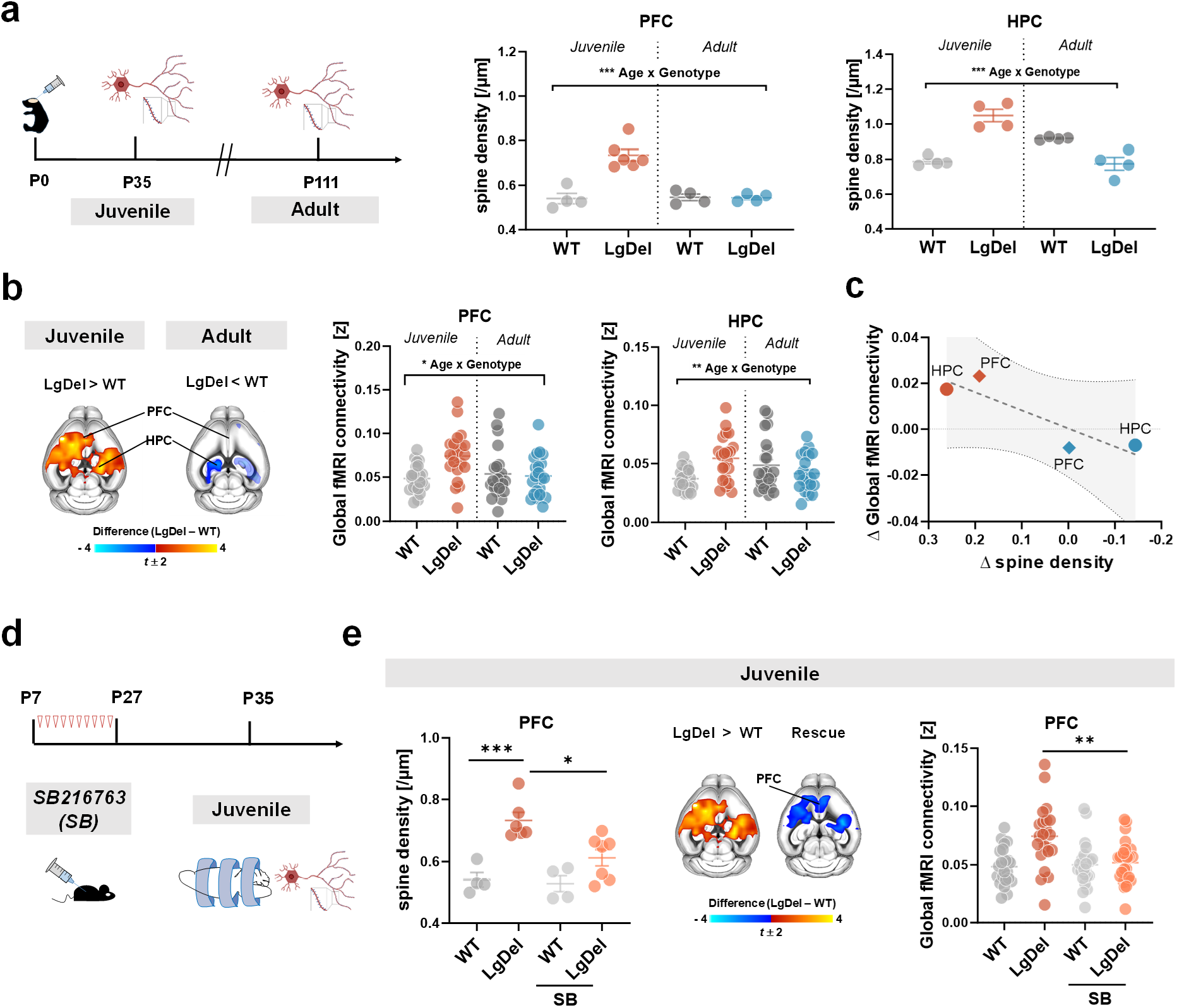
Developmental fMRI dysconnectivity is paralleled by GSK3β-dependent alterations in dendritic spine density. (a) Dendritic spine density measurements across development in the prefrontal cortex (PFC) and the hippocampus (HPC) revealed a significant age x genotype interaction. (b) Voxel-wise global fMRI connectivity in these two areas exhibited a similar developmental trajectory, with a significant age x genotype interaction both in PFC and HPC. (c) Intergroup differences in synaptic density and fMRI global connectivity show a quasi-linear relationship (R^2^ = 0.8). (d) Experimental timeline of the GSK3β inhibition treatment protocol (from p7 to p27), followed by fMRI and spine density measurements. (e) Developmental GSK3β inhibition rescued spine density increase in PFC of juvenile mice. Similarly, the same treatment restored global fMRI connectivity alterations in juvenile LgDel mice. Rescue map represents the comparison between LgDel SB-treated mice (LgDel SB) with LgDel vehicle-treated mice (LgDel) (|t| > 2.0, FWER cluster-corrected). Regional quantification of global fMRI connectivity across 4 groups (WT, WT SB, LgDel, LgDel SB) confirmed this result. Errors bars represents SEM. *p<0.05, **p < 0.01, ***p < 0.001. [HPC, hippocampus, PFC, prefrontal cortex].

To putatively link alterations in spine density to the observed fMRI dysconnectivity, we next quantified global fMRI connectivity in PFC and hippocampus, and compared the observed fMRI connectivity differences with the corresponding alterations in spine density (Figure 3b). Interestingly, global fMRI connectivity quantifications broadly recapitulated the group-and age-specific profile observed in dendritic spine count experiments. Specifically, fMRI connectivity was only transiently increased in the PFC of LgDel mice at the juvenile stage, while juvenile fMRI hyperconnectivity in the hippocampus fully reverted to hypoconnectivity in adulthood. Moreover, inter-group differences in global fMRI connectivity and spine density were found to covary (Figure 3c). These findings suggest that developmental hyper-and hypoconnectivity in LgDel mice might reflect increased and decreased synaptic density, respectively.

If alterations in synaptic density are causally implicated in the fMRI dysconnectivity we observed in LgDel mice, normalization of synaptic pathology should be associated with a normalization of fMRI phenotypes. To test this hypothesis, we leveraged previous reports that the GSK3β pathway is hyper-active in rodent models of 22q11DS ^40^. GSK3β is a critical regulator of synaptic development and function ^40, 41, 42^. GSK3β overactivity may thus lead to impaired synaptic homeostasis and interfere with the large-scale synaptic pruning and refinement that occurs during childhood and adolescence, resulting in altered dendritic spine density.

To test the hypothesis the GSK3β-hyperactivity may underlie synaptic alterations observed in LgDel mice, we pharmacologically inhibited GSK3β via administration of the selective antagonist SB-216763 (SB) during the first three post-natal weeks (Figure 3d). This developmental period is characterized by robust spine growth and synaptogenesis, as well as high expression of GSK3β ^42, 43^. We next measured dendritic spine density in the PFC and whole brain fMRI connectivity in two separate groups of mice (Figure 3e, Figure S5).

Because SB was administered during an early developmental window, we expected this treatment to be able to (at least transiently) normalize the synaptic and fMRI connectivity alterations we found in juvenile mice. Corroborating this notion, developmental GSK3β inhibition significantly reduced spine density surplus in the PFC of juvenile LgDel mice (Figure 3e, two-way ANOVA, *Sidak’s multiple comparisons test*, LgDel SB vs LgDel, p = 0.01, Figure S5). This treatment also robustly normalized fMRI hyperconnectivity in mice imaged at the same prepubertal stage, as probed either with global fMRI mapping or seed-based mapping of the default mode network (DMN; two-way ANOVA, *Sidak’s multiple comparisons test*, LgDel SB vs LgDel, p = 0.0009, Fig 3e, Fig S6). Confirming the specificity of these findings (and corroborating a possible link between synaptic alterations and fMRI dysconnectivity) dendritic spine density and global fMRI connectivity were found to be similarly unaltered in the PFC of adult mice receiving SB or vehicle administration (Figures S6 and S7). Taken together, these results implicate GSK3β-dependent synaptic-alterations in the establishment of developmental fMRI dysconnectivity we identified in LgDel mice. The observed normalization of fMRI dysconnectivity connectivity was transient, and did not extend to adulthood, as we found that foci of global hippocampal fMRI hypoconnectivity in adult LgDel were not affected by postnatal pharmacological treatment with SB (Figure S7).

To probe the behavioral relevance of these connectivity alterations we next assessed social behavior and cognitive function in juvenile and adult mice, with and without treatment with SB. These investigations highlighted the presence of translationally relevant behavioral alterations in LgDel mice. At the juvenile stage, LgDel mice displayed impaired sociability in a social habituation-dishabituation test (two-way ANOVA, LgDel vs. WT *p* = 0.04, Figure S8). However, these impairments were unchanged by GSK3β inhibition (LgDel SB vs. LgDel *p* = 0.07). Juvenile LgDel mice did not show overt alterations in social reward processing, as probed with a social conditioning place preference task (LgDel vs. WT *p* = 0.26, Figure S8). This behavior was also not affected by GSK3β treatment (LgDel SB vs. LgDel *p* = 0.97).

Similar to what was observed at the juvenile stage, adult LgDel mice displayed reduced sociability (two-way ANOVA, LgDel vs. WT *p* = 0.047), but broadly preserved social memory (LgDel vs. WT *p* = 0.18, Figure S8) in a three-chamber social approach task. Pharmacological GSK3β inhibition did not mitigate these socio-behavioral alterations. Finally, adult LgDel mice exhibited impaired cognition in a temporal order memory task (two-way ANOVA, LgDel vs WT, *p* = 0.002). In contrast to the other paradigms, this deficit was ameliorated by GSK3β inhibition (LgDel SB vs LgDel, *p* = 0.003, Figure S8). Taken together, these investigations provide evidence of social alterations and impaired memory-related cognition in adult LgDel mice, with only the latter being improved by GSK3β inhibition. These observations also suggest that, while modulating GSK3β activity may re-establish (synaptic and fMRI) brain connectivity and memory-related cognition in LgDel mice, this pathway may be less implicated in the control of social behavior.

To more rigorously probe the relationship between behavioral alterations and fMRI dysconnectivity in LgDel mice, we used partial least square analysis (PLS, ^44^) to model covariance between fMRI connectivity and behavioral performance in the tests described above. By using a multivariate approach, we sought to control for statistical bias related to repeated univariate testing of multiple behavior ^45^. This “brain-behavior” relationship was assessed for connectivity in the DMN and hippocampal networks, i.e., two systems encompassing key substrates for social and memory-related functions ^10, 46, 47^. We also assessed possible modulation by GSK3β-inhibition of brain-behavioral covariance patterns by entering treatment as a binary design variable in the behavioral matrix.

In line with previous reports ^10, 47^ we found, both in juvenile and adult wild-type mice, significant covariance between sociability and fMRI connectivity in striatocortical components of the DMN (Figure S9). Interestingly, in LgDel mice this relationship was disrupted, such that neither prefrontal nor hippocampal patterns of connectivity were found to covary with behavioral profiles in the same fashion as wild-type mice (Figure S9). The lack of significant covariance between treatment and behavioral performance further suggested that the brain-behavior relationship was not restored by pharmacological GSK3β inhibition. These results suggest that connectivity-behavior relationships are disrupted in LgDel mice, and that this relationship is not restored by developmental treatment with SB.

### Developmental dysconnectivity in individuals with 22q11DS occurs in brain regions enriched for synaptic genes interacting with GSK3β

Our mouse investigations suggest that synaptic-related mechanisms, and specifically GSK3β pathway dysregulation, may underlie fMRI dysconnectivity in LgDel mice. To probe the generalizability of these findings to human 22q11DS, we used a gene decoding approach to relate the observed developmental dysconnectivity to spatial patterns of post-mortem gene expression in the same regions ^48^. We thus identified a list of *N* = 3897 transcripts (henceforth “decoded genes”) that are spatially enriched in areas exhibiting developmental dysconnectivity in individuals with 22q11DS (Figure 4a). We next ran an enrichment analysis for genes implicated in synaptic signaling as defined by gene ontology (GO) categories ^49^. In keeping with our hypothesis, we found a significant enrichment for multiple synaptic-related GO gene lists (Figure 4b). Interestingly, no enrichment for glial-, microglia-or immune-related genes was observed.

**Figure 4.**
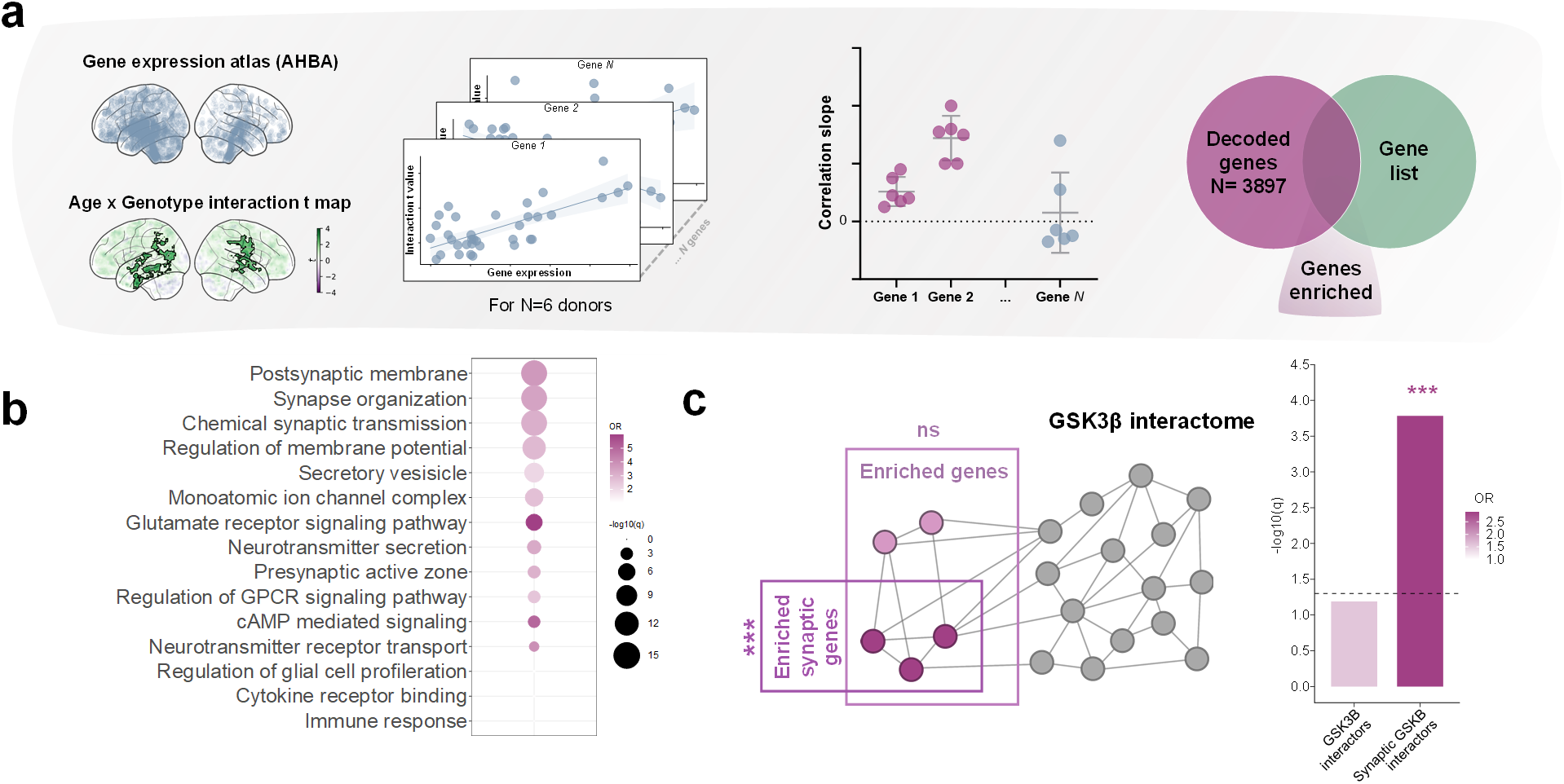
Gene decoding supports involvement of synaptic mechanism in 22q11DS developmental dysconnectivity. (a) Illustration of gene decoding and gene enrichment analyses used to investigate molecular mechanisms underlying developmental reconfiguration. The term “decoded genes” refers to genes that were spatially enriched (i.e. genes that displayed significantly higher expression) in areas that undergo hyper-to-hypoconnectivity reconfiguration in 22q11DS. (b) Decoded genes are specifically and significantly enriched for synaptic-related GO gene lists, corroborating the involvement of synaptic mechanisms in 22q11DS dysconnectivity. Color scale indicates odds ratio, while size of the dots represents –log_10_(q-value). Only visible dots were statistically significant at q < 0.05. (c) Schematic representation of gene enrichment analyses (left) and results (right) showing significant enrichment for synaptic interactors of GSK3β. Bar color indicates odds ratio, length represents –log_10_(q-value). Horizontal dashed line (in grey) represents significance at q < 0.05. * p < 0.05, ** p < 0.01, *** p < 0.001.

To corroborate putative mechanistic correspondences with our mouse data, we next investigated whether brain regions undergoing developmental dysconnectivity in individuals with 22q11DS would be specifically enriched for GSK3β -related synaptic genes. To test this hypothesis, we first examined whether the decoded genes would be enriched for genes that interact at the protein level with GSK3β. This analysis revealed a weak enrichment of GSK3β interactors in the decoded gene list (OR = 1.57, *q* = 0.06, overlapping genes N = 129, Figure 4c). However, when we pruned this list to solely retain GSK3β interactors involved in synaptic processes as per the curated SYNGO database ^50^, we found the decoded list to be robustly enriched for these transcripts (OR = 2.91, *q* = 0.0002, overlapping genes N = 45, Figure 4c). To test the spatial specificity of this result, we applied gene decoding on k = 1000 spatial autocorrelation-preserving surrogate maps. Enrichment analyses of the gene lists decoded from these surrogates suggest that our enrichment was unique to our developmental dysconnectivity pattern, as the size of our enrichment mapped above 99^th^ percentile of the obtained null distribution. These results suggest that GSK3β-related synaptic mechanisms may also underlie developmental dysconnectivity in human 22q11DS dysconnectivity.

### Developmental dysconnectivity within sensory-motor systems predicts social impairment in 22q11DS

To investigate the potential clinical relevance of the observed developmental dysconnectivity in 22q11DS, we probed whether the decoded gene list would be significantly enriched for subcategories of genes known to be involved in neurodevelopmental disorders. We found that the pattern of developmental dysconnectivity we identified in 22q11DS individuals (Figure 2b) was spatially enriched for genes involved in schizophrenia as well as for genes associated with autism. However, enrichment for autism-relevant gene lists, especially genes whose expression is reduced in post-mortem cortical samples of autistic people, was remarkably stronger and exhibited especially high significance (^51^, OR = 4.15, *q* < 0.0001, overlapping genes N = 917; SFARI gene list, OR = 2.08, *q* < 0.0001, overlapping genes N =334, Figure 5a). No enrichment for genes associated with bipolar disorder^52^ or ADHD ^53^ was found (Figure 5a).

**Figure 5.**
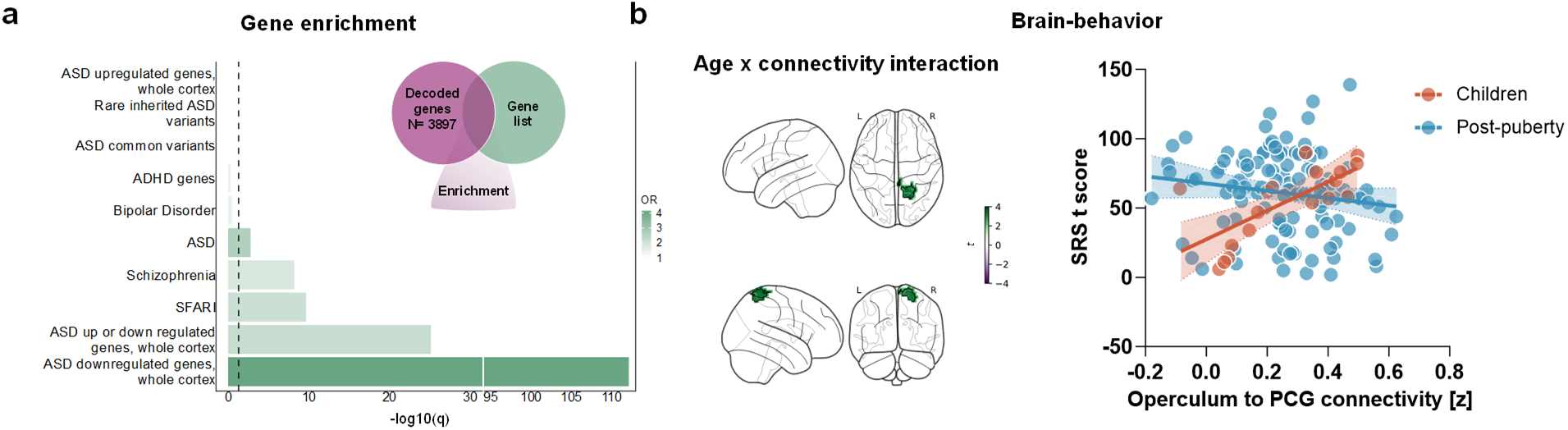
Developmental fMRI dysconnectivity within somatomotor areas predicts reciprocal social behavior in 22q11DS. (a) Gene enrichment analyses showing highly significant enrichment between decoded genes and schizophrenia-and ASD-related genes. Bar color indicates odds ratio, length represents –log_10_(q-value). Vertical dashed line (in grey) represents significance at q < 0.05. (b) Voxelwise (left of panel) and ROI-based quantifications of relationship between opercular cortex connectivity and SRS score in 22q11 deletion carriers. Map represents areas of significant age x connectivity interaction in predicting SRS score (|t| > 2.0, FWER cluster-corrected). Quantification of relationship between opercular cortex to postcentral gyrus (PCG) connectivity and SRS within the two age groups separately (right).

The observation of a disproportionately high enrichment for genes exhibiting functional downregulation in autism (Figure 5a, p = 10^-113^) led us to hypothesize that the observed developmental dysconnectivity may be associated with autism-relevant symptoms in people with 22q11DS. Such notion would be consistent with the prepubertal manifestation of hyperconnectivity in 22q11DS individuals. We thus selected a sub-set of N = 119 22q11.2 deletion carriers (N = 18 children, N = 111 adults) for which we had corresponding Social Responsiveness Score (SRS) data and used a linear model to test the hypothesis that magnitude of fMRI dysconnectivity in areas undergoing the developmental switch (i.e., those showing significant age x genotype interaction), would be predictive of reciprocal social behavior impairment, assessed via SRS score. If this is the case, SRS scores in 22q11DS individuals would be positively associated with fMRI hypoconnectivity in childhood, and negatively associated with fMRI hypoconnectivity in adulthood, respectively.

The application of this model to fMRI global connectivity maps did not reveal such an association, either in voxelwise mapping (p > 0.05, FWER cluster-corrected), or within dysconnectivity clusters (i.e., those mapped in Figure 2b, S2, p > 0.12, all areas and conditions). However, global fMRI connectivity is an aggregate voxel-wise measure that incorporates multiple, contrasting network-specific associations. We thus applied our model to individual somatosensory networks that we previously observed to undergo developmental reconfiguration via seed-based mapping. Linear modelling of the somatomotor/auditory network (see Figure 2c) revealed a cortical cluster exhibiting a developmental interaction between SRS and fMRI connectivity (age x fMRI connectivity, β = 106.3, p=0.009, Figure 5b). As hypothesized, this relationship was characterized by a positive association between fMRI connectivity and SRS in childhood (β = 103.8, p =0.003) which reverted to a (weaker) negative association after puberty (β = -26.21, p = 0.11). These results suggest that the hyper-to-hypoconnectivity switch in sensory-motor brain regions in 22q11.2 deletion carriers is predictive of autism-relevant traits.

## Discussion

In the present work we describe a cross-species investigation of the neural determinants and developmental course of functional connectivity in 22q11DS, a highly penetrant genetic risk model for schizophrenia. We report, for the first time, converging cross-species evidence of developmental dysconnectivity in 22q11DS, with fMRI hyperconnectivity reverting to hypoconnectivity over puberty, both in LgDel mice and in people harboring a chromosomal 22q11.2 deletion. In the mouse, fMRI dysconnectivity is accompanied by GSK3β-dependent spine density alterations mirroring functional connectivity differences. In keeping with rodent findings, brain regions exhibiting developmental reconfiguration in human 22q11DS are enriched for synaptic-related genes that interact with GSK3β. Lastly, we report that functional dysconnectivity in somatomotor components of this network is predictive of social alterations in 22q11.2 deletion carriers. Collectively, our results provide evidence of synaptic-related developmental dysconnectivity in 22q11DS.

Resting state fMRI connectivity mapping has been previously used to describe network correlates of pathology in 22q11DS. Evidence of hypoconnectivity in hippocampal and key nodes of the default mode network (DMN) has been consistently reported in previous investigations of fMRI connectivity in individuals with 22q11DS ^23, 54, 55^. Initial studies investigating the developmental evolution of fMRI dysconnectivity in 22q11DS have also been reported, although with caveats related to the limited availability of fMRI investigations in children with 22q11DS. These investigations have shown that the developmental trajectory of fMRI connectivity within the DMN, hippocampus and in thalamo-cortical areas may be atypical in individuals with 22q11DS ^54, 56^. Evidence of possible developmental alteration in functional connectome organization in 22q11DS has also been reported across development ^57^. Our study expands previous investigations in two main directions. First, by integrating human studies with controlled, parallel investigation in a rodent model, we were able to generate testable hypotheses regarding the developmental trajectory of fMRI dysconnectivity in 22q11DS with a degree of confidence currently unattainable in clinical research ^13^. Rodent investigations were instrumental in the identification of a critical developmental window during which cortical dysconnectivity dramatically reconfigures in 22q11DS. Second, our cross-species work allowed to probe the mechanisms underlying these findings. From a methodological standpoint, our results epitomize the potential of cross-species research in biologically decoding the mechanism of fMRI dysconnectivity in humans.

By longitudinally mapping fMRI connectivity in LgDel mice, we provide a detailed account of developmental dysconnectivity in an established rodent model of 22q11DS ^20^. The presence of reduced hippocampal connectivity in adult LgDel mice is in agreement with the results of fMRI studies in other rodent models of 22q11DS ^58, 59^, and recapitulates functional neuroimaging findings that have been documented in individuals with 22q11DS ^23, 54^. Extending these previous investigations, our work shows that this fMRI hypoconnectivity develops over puberty, is preceded by widespread cortical hyperconnectivity and it is plausibly mediated by synaptic-dependent mechanisms.

The presence of altered synaptic alterations is a feature shared by multiple transgenic mouse models of 22q11DS, where reduced synaptic density in hippocampal regions has been typically observed ^35, 37, 60^. We report that in cortical and hippocampal regions dendritic spine alterations are prominent, and dramatically evolve over puberty. The presence of supranumeral synapses and fMRI hyperconnectivity in cortical and hippocampal regions in prepubertal LgDel animals is broadly consistent with recent work linking synaptic surplus to aberrant fMRI coupling in a mouse model of syndromic autism ^10^. This finding is also consistent with our observation of aberrant extension of somatomotor connectivity networks in children with 22q11DS, (Figure 2c) where it could indicate altered synaptic refinement and impaired network maturation ^61, 62^. Overall, the emerging model is one in which deficient synaptic pruning during early development is followed by compensatory (PFC) or overcompensatory (hippocampus) synaptic elimination over puberty. The biological mechanisms underlying aberrant synaptic elimination in hippocampal regions were not investigated here. However, the involvement of aberrant microglia-related signaling and synaptic elimination represents a possible explanation for this remodeling. In keeping with this notion, we recently found evidence of increased microglial complement systems C4 expression in adolescent LgDel mice ^63^ a phenotype that has been associated with aberrant synaptic pruning over puberty ^39^.

While the mechanisms leading to dendritic synaptic surplus in juvenile mice may be manifold, our pharmacological manipulations suggest that GSK3β inhibition is sufficient to prevent this phenotype in LgDel mice. Regional enrichment for GSK3β synaptic interactors in areas of human 22q11DS neural dysconnectivity corroborates the relevance of this finding for human 22q11.2 deletion carriers. Importantly, hyperactivity of GSK3β is etiologically linked to 22q11DS via Zdhhc8 haploinsufficiency, a hemizygously deleted genes in the 22q11.2 locus ^40^. From a mechanistic standpoint, these results are consistent with GSK3β’s pivotal role in controlling synaptic development, function and homeostasis ^41^. GSK3β hyperactivity may thus influence synaptic homeostasis via an indirect modulation of plasticity mechanisms underlying synapse formation and elimination ^64^ or by interfering with autophagic pathways essential for synapse maintenance ^10, 65^. Recent evidence also points at a crucial role of microglia in mediating the phagocytosis of synaptic debris during pruning ^66, 67^, a pathway in which GSK3β is also implicated ^68^. While our data plausibly link aberrant GSK3β-signalling to juvenile fMRI hyperconnectivity, they do not conclusively establish whether fMRI hypoconnectivity is similarly GSK3β-dependent. The fact that this post-pubertal phenotype was not rescued by developmental SB administration may simply reflect a time-limited action of the drug, or the involvement of GSK3β-independent synaptic elimination. In this respect, a contribution of microglial mechanisms or other known 22q11.2 synaptic interactors (i.e. COMT, DGCR8, RTN4R, and SEPT5) cannot be ruled out.

The observation of peri-pubertal dysconnectivity remodeling in LgDel mice, and its confirmation in a large clinical cohort is of interest in the light of the high risk 22q11DS confers for both early and late onset neurodevelopmental disorders ^14^. While evidence of atypical developmental dysconnectivity has been reported in hippocampal ^54^ or thalamocortical areas ^25^, to the best of our knowledge our work is the first to detail widespread cortical hyperconnectivity in prepubertal 22q11DS individuals, a finding associated with aberrant anatomical extension of somatomotor-network connectivity. The parallel identification of this phenotype in mice and human increases our confidence in this finding, despite the still limited number of fMRI scans available in prepubertal 22q11DS individuals. While the clinical relevance of this connectivity reconfiguration remains to be fully determined, our analyses revealed that dysconnectivity in sensory-motor areas was predictive of autism-related traits, specifically impairments in reciprocal social behavior. This notion is consistent with the observation of a disproportionately high enrichment of autism-relevant genes in the same regions undergoing developmental switching, and with the developmental conceptualization of autism (and autism-related disorders) as dysconnectivity syndromes characterized by hyper-connectivity in childhood, and hypoconnectivity in adult stages ^69^. It should also be noted that fMRI connectivity in somatomotor areas has been recently found to be associated with symptom severity in idiopathic autism ^70^. We however acknowledge that, albeit plausible and enticing, this association does not entail causality. Further proof of the specificity and replicability of this brain-behavior association will require extensions to larger patient cohorts, and cross-referencing to other symptoms domains (e.g., psychosis-related traits), an analysis that was not feasible in the present study owing to the paucity of both brain and behavioral data in children with 22q11DS.

The reduced sociability we observed in LgDel mice recapitulates social impairments described in individuals with 22q11DS ^71, 72^. However, brain-behavior investigations in mice did not appear to directly align with corresponding human results. Specifically, as previously reported ^10^, we found that in wild-type mice connectivity within the DMN and its striatal components are positively correlated with social behavioral performance. However, this relationship was disrupted, and as such uncorrelated with social behavior, in LgDel mice. This finding is not entirely surprising, owing to the evolutionary distance, and the highly diverging substrate underlying higher order socio-cognitive functions in rodents and humans. Indeed, in rodents social processing is strongly rooted in olfactory encoding, while in humans it involves processing of visual functions that are poorly developed in mice ^73, 74^.

Of note, adult LgDel mice also exhibited cognitive dysfunction in the temporal order memory domain, consistent with results in other mouse models of 22q11DS ^75^, and the observation of hippocampal-dependent memory alterations in humans ^76^. Interestingly, developmental inhibition of GSK3β did not ameliorate social dysfunction at either age, but was effective in reversing memory impairment in adult mice, as previously documented ^77^. The clear dissociation between early and late effect of GSK3β inhibition on connectivity and cognitive deficits suggests that fMRI connectivity reflects basic coupling mechanisms (e.g., dendritic spine integration ^10^) that are not necessarily directly related or relevant to higher order cognitive functions related to brain plasticity, like temporal order memory. As indicated by a growing body of evidence, synaptic plasticity can instead be profoundly impaired by GSK3β overactivity, resulting in aberrant LTP-like activity ^42^. GSK3β overactivity can also influence control of transcriptional and metabolic processes ^64, 78, 79^. Future investigations are required to disambiguate these contributions and relate them to specific symptoms and manifestations of 22q11DS.

In conclusion, our work provides the first cross-species evidence of fMRI dysconnectivity in 22q11DS over puberty. Mouse and human studies converge to suggest that the observed fMRI alterations are underpinned by synaptic-dependent mechanisms, encompassing a pivotal involvement of GSK3β signaling. This developmental dysconnectivity is also predictive of social alterations in human 22q11.2 deletion carriers. Taken together, our results shed light on the etiopathological underpinnings of fMRI dysconnectivity 22q11DS.

## Acknowledgments

The authors would like to dedicate this work to the cherished memory of Andrea Baracchino. This work was supported by a MCHRI Uytengsu-Hamilton 22q11 Neuropsychiatry Research Award (UH22QEXTFY22-04), the NIH (1R21MH116473-01A1 to A. Gozzi) and the European Research Council (ERC—DISCONN, No. 802371). A. Gozzi was also supported by Brain and Behavior Foundation 2017 (NARSAD—National Alliance for Research on Schizophrenia and Depression), the Telethon foundation (GGP19177) and by an endowment by Paolo and Sara Baracchino. M. Pagani was supported by the European Union’s Horizon 2020 research and innovation programme under grant agreement No. 845065 (Marie Sklodowska-Curie Global Fellowship - CANSAS). M.V. Lombardo acknowledges funding by the European Research Council (ERC) under the European Union’s Horizon 2020 research and innovation programme under grant agreement No 755816.

## Materials and Methods

### Mouse studies

#### Ethical statement

Animal studies were conducted in accordance with the Italian Law (DL 26/2014, EU 63/2010, Ministero della Sanità, Roma) and the recommendations in the National Institutes of Health Guide for the care and use of laboratory animals. Animal research protocols were reviewed and consented to by the animal care committee of the Istituto Italiano di Tecnologia and the Italian Ministry of Health specifically approved the study protocol (authorization no. 752/19 to A. Gozzi).

#### Animals and experimental cohorts

Animals were housed under controlled temperature (21 ± 1°C) and humidity (60 ± 10%) and maintained on a 12-hour light/dark cycle, with food and water available ad libitum. LgDel mice ((Del(16Dgcr2-Hira)1Rak, Jackson laboratories) were backcrossed for more than ten generations with C57BL/6J mice and genotyped as previously described ^27^. “Wild-type” (WT) littermates were used as control, WT mice.

Mouse fMRI and behavioral investigations encompassed a cross-sectional treatment protocol with four cohorts of age matched mice (mixed sexes): WT mice treated with SB216763 (2 mg/kg i.p.; WT SB, n = 22; n = 12 males, n = 10 females); WT mice treated with vehicle (2mg/ml DMSO in polyethylene glycol 400; WT, n = 22; n = 14 males, n = 8 females); LgDel mice treated with SB216763 (2 mg/kg i.p., ^77^; LgDel SB, n = 23; n = 14 males, n = 9 females); LgDel mice treated with vehicle (LgDel, n=21; n = 12 males, n = 9 females). SB216763 or vehicle were administered every other day from P7 to P27 at 9 a.m as previously described ^77^. The drug was initially dissolved in dimethyl sulfoxide (DMSO) at a concentration of 20 mg/ml. On the days of injection, drug stock solutions were freshly diluted with polyethylene glycol 400 (PEG400) to achieve a final concentration of 2 mg/kg as in Tamura et al. (2016).

Mice underwent two sessions of fMRI imaging, the first at the juvenile stage (p35 ± 1), and the second after puberty (p109 ± 3). Behavioral assessments of sociability were conducted on the same mice at the age of 30 ± 2 days, immediately before resting-state functional MRI (rsfMRI) imaging. Behavioral tasks in adulthood were conducted after the second imaging session. Prior to the second imaging session two mice from the LgDel SB group exhibited health issues and were therefore removed from the study, resulting in a final sample size of n = 23. All behavioral testing and procedures were conducted during the light phase of the cycle. Experimenters were blind to mouse treatments during testing and behavioral scoring.

#### Resting state fMRI acquisition

For resting state functional MRI mice were anesthetized with isoflurane (5% induction), intubated, and artificially ventilated (2% maintenance) ^8, 10^. After surgery, isoflurane was discontinued and replaced with halothane (0.8%). Functional data acquisition started 30 min after isoflurane cessation. Ventilation parameters were adjusted to maintain normo-physiological p_a_CO_2_ (<40 mmHg) and p_a_O_2_ levels (>90 mmHg, corresponding to >98% hemoglobin saturation). fMRI images were acquired a 7T MRI scanner (Bruker Biospin, Milan) with Bruker Paravision software (v6) using a 72-mm birdcage transmit coil and a 4-channel solenoid coil for signal reception ^29^. Single-shot BOLD rsfMRI time series were acquired using an EPI sequence with the following parameters: TR/TE 1000/15 ms, flip angle 60°, matrix 98 × 98, FOV 2.3 × 2.3 cm, 18 coronal slices, slices thickness 550 µm for 1920 volumes.

#### fMRI connectivity analysis

rsfMRI images were preprocessed as previously described ^10^. Briefly, the initial 50 volumes of the time series were removed to allow for T1 and gradient thermal equilibration effects. Data were next despiked, motion-corrected, skull-stripped and registered to a BOLD reference template. Motion traces of head realignment parameters (three rotations and translations) and mean ventricular signal were used as nuisance covariates and regressed out. All fMRI time series were next band-pass filtered (0.01-0.1 Hz) and spatially smoothed with a full width at half maximum of 0.6 mm. Finally, we carried out censoring of BOLD volumes flagged for high motion using a framewise displacement threshold greater that 0.05 mm).

To identify “hotspots” of dysconnectivity in a spatially-unbiased fashion, we computed global functional connectivity at the voxel level. This metric quantifies the temporal correlation between fMRI signal of a voxel with that of all the other voxels in the brain. Global connectivity is predominantly skewed toward long-range connections, defined here as those exceeding 600 µm from a given voxel, which account for 99.77% of all brain voxels ^47^. Pearson’s correlation scores were first transformed to z scores using Fisher’s r to z transformation and then averaged to provide the final connectivity scores. Functional connectivity network mapping was carried out using a seed-based analysis by placing predefined seeds (3 x 3 x 1, voxels) in regions of interest selected based on our global connectivity results (Supplementary Figure 1). Between-group differences were mapped using a two-tailed Student’s *t* test (|t| >2, p< 0.05) and family-wise error (FWE) cluster corrected for multiple comparisons using a cluster defining threshold of P< 0.05 as implemented in FSL. Voxel-wise interaction between Genotype and Age factors was assessed using a linear mixed effects model (lmer4 package in R) with within-subject variability as random intercept (+1|subject). The obtained t score maps were (FWE) cluster-corrected using a cluster threshold of p=0.05.

To depict effect size of mapped fMRI differences, z-scored fMRI connectivity from cubic volumes of interest (VOIs) (5 x 5 x 1) were extracted and plotted (Supplementary Figure 1). The resulting values were tested for statistical significance using a linear mixed-effects analysis. Brain regions showing connectivity differences in previous age x genotype interaction analysis (Fig. 1e-f) were further investigated for the presence of sex x genotype interaction at both ages. These analyses did not provide evidence of any statistically significant sex x genotype interaction.

#### Dendritic spine quantification

To highlight dendritic spines of cortical and hippocampal pyramidal neurons for postmortem quantifications, 2µl of AAV8-hSyn-GFP 10^11^ IU/ml were bilaterally injected in lateral ventricles of newborn WT and LgDel mice at postnatal day 1. Four mice cohorts were analyzed at each developmental stage: PFC counts in juvenile mice, WT Veh n=4 (n = 2 males; n = 2 females), WT SB n = 4 (n = 3 males; n = 1 female), LgDel Veh n = 6 (n = 1 male; n = 5 females), LgDel SB n =7 (n = 4 males; n = 3 females); PFC counts in adult mice, WT Veh n = 4 (n = 3 males; n = 1 female), WT SB n = 4 (n = 2 males; n = 2 females), LgDel Veh n = 4 (n = 2 males; n = 2 females), LgDel SB n = 4 (n = 2 males; n = 2 females). Spine counting in HPC was conducted in the same animals as for PFC. In juvenile mice hippocampal spines were counted in 4 animals (n = 1 male; n = 3 females). Vehicle or SB administration was carried out from p7 to p27 as described above. At postnatal days p33 (juvenile) or p112 (adult), mice were sacrificed via transcardial 4% paraformaldehyde PBS-perfusion. Brains were sectioned with a vibratome (Leica) at 50µm (coronal) and processed for free-floating immunofluorescence. Briefly, tissue was incubated in blocking solution (5% horse serum in PBTriton 0.5%) 1h at RT and subsequently incubated overnight at 4°C in primary antibody solution using Rabbit anti-Green-Fluorescent-Protein (GFP; 1:1000 ThermoFisher Scientific, A6455). The following day sections were rinsed three times in PBTriton 0.5% and incubated overnight at 4°C in secondary antibody solution using Goat anti-Rabbit-Oregon Green 488 (1:500 Invitrogen, 011038). Next, samples were rinsed in PBTriton, counterstained with DAPI (1:1000) and mounted with AquaPolymount (Polyscience). High-resolution images dendritic spines were acquired using a Nikon A1 Eclipse confocal microscope equipped with NIS element AR v4.20.03 software (Nikon) and a 60x plan-apo oil immersion objective with a 2x digital magnification (1024x1024px, 0.1µm/px). Spine density quantification was performed on basal dendrites of layer V prefrontal cortex pyramidal neurons (PFC, Bregma from 2.58 to 2.34 mm) and on apical dendrites of CA1 hippocampal neurons (HPC, Bregma from -2.92 to -3.40). Spine counting was performed using the Filament function of Imaris-Microscopy Image Analysis Software by Bitplane (v7.2.3). A single first order dendrite per neuron was analyzed for a total of 10-15 neurons for each animal. The analysis was performed by an operator blind to the genotype. Data are expressed as spine number per µm of dendrite length. Statistical analysis was performed using Prism GraphPad (v9.2).

### Behavioral Testing

#### Habituation/dishabituation social interaction test

Juvenile mice were tested as previously reported ^80^ in lightly illuminated (5±1 lux) GR900 Tecniplast cages (904 cm^2^). Mice were placed individually in the testing cage 1 hour prior to the test. A stimulus mouse of the same sex, strain, and age was introduced into the testing cage for a 1-minute interaction. After the 1-minute trial, the stimulus mouse was removed for 3 minutes and placed back in its home cage. This procedure was repeated four times. Subsequently, a fifth 1-minute dishabituation trial was conducted, introducing a new stimulus mouse to the testing cage. The time spent interacting (interaction time), which included the sum of nose-to-nose sniffing, anogenital sniffing, and following, was recorded across all trials. The interaction time across the first four trials was summed up and analyzed using a two-way ANOVA.

#### Social conditioning place preference (SCPP) test

Juvenile mice also underwent a social conditioning place preference (SCPP) test, adapted from a previous study of ^81^. The test was conducted in a three-chamber arena, with each chamber measuring 20x40x22 cm. Two different environments were used for the conditioning sessions, with each environment represented by different types of bedding: corncob bedding (SCOBIS DUE G8, Mucedola) and aspen bedding (SCOBIS QUATTRO, Mucedola). To prevent any bias toward one environment from affecting the results, a 30-minute pre-conditioning test was conducted on the first day. In this test, both environments were randomly allocated to each of the two external chambers. To mitigate potential natural biases that might obscure the effects of social conditioning, the less preferred context was paired with social enrichment. Mice that displayed a pre-conditioning preference of over 70% for one environment type were excluded from subsequent conditioning. This led to the exclusion of one WT mouse, two LgDel mice, and three LgDel SB mice, resulting in a final sample size of WT n=22, LgDel n=21, WT SB n=22, and LgDel SB n=22.

After the pre-conditioning phase, mice underwent a 24-hour social conditioning session with their cagemates in the environment where they initially spent less time. Twenty-four hours later, they were placed in the other environment type without their cagemates. The next day, a 30-minute post-conditioning session was conducted following the same procedure as the pre-conditioning session. The performance in the pre-conditioning and post-conditioning sessions within the same group was assessed through a paired Student’s t-test. To address potential variations in locomotory behavior within the apparatus and baseline preferences, comparisons between experimental conditions were made using a preference score. This score was calculated as the percentage of time spent in the social context, normalized by the total time in compartments, with baseline values subtracted as described in ^82^. Data were analyzed using a 2-way ANOVA with genotype and treatment factors followed by a Sidak’s multiple comparisons test.

#### Three-chambered social approach tasks

After the second imaging session, adult mice were tested in a three chamber test, following the protocol described in ^83^. The apparatus consisted of a three-chamber box, with each chamber measuring 20x40x22 cm, featuring two sliding doors (5x8 cm) that opened into the central compartment, and two wire cages (15 cm in height) where a ‘stimulus’ mouse was introduced. The ‘stimulus’ mouse was an experimentally naïve mouse, with no prior interactions with the testing mouse. These stimulus mice were of the same sex, background, aged between 2-6 months, and approximately matched in body weight to the testing mouse (within 5 grams). Prior to the task, the stimulus mice were habituated to the wire cups four days in advance to minimize signs of agitation and aggression.

The task commenced with a 10-minute habituation phase in the central compartment, followed by an additional 10-minute habituation phase encompassing all three chambers. The ‘sociability phase’ (10 minutes) began immediately after the habituation phase. During this phase, the testing mouse was introduced into the central zone and allowed to explore side compartments, one containing an empty cup and the other containing a cup with a novel stimulus mouse. The locations of the empty and ‘mouse’ cups were randomly assigned. A ‘sociability index’ was calculated as the difference between the time spent sniffing the mouse cup and the time spent sniffing the empty cup, normalized by the total time spent sniffing. Subsequently, a ‘social novelty phase’ (10 minutes) was conducted, with a new stimulus mouse introduced in the empty cup. A ‘social novelty index’ was calculated as the difference between the time spent sniffing the cup with the new stimulus mouse compared to the cup with the old stimulus mouse, normalized by the total time spent sniffing.

One hour later, a ‘social memory phase’ (10 minutes) was carried out. A ‘social memory sniffing’ was calculated as the difference between the time spent sniffing the cup with the newest stimulus mouse and the cup hosting the stimulus mouse from the sociability phase, normalized by the total time spent sniffing. In this phase, the same stimulus mouse from the sociability phase was placed in the same side as before, while a novel stimulus mouse was placed in the other side. Two LgDel animals were exempted from testing due to health issues (corneal opacity) resulting on a final sample of WT n=21, LgDel n=20, WT SB n=22, and LgDel SB n=22. Data were analyzed using a 2-way ANOVA with genotype and treatment factors followed by a Sidak’s multiple comparisons test.

#### Temporal order memory task

Adult mice also underwent a temporal order memory task ^84^. This task assesses recency discrimination by measuring the time spent exploring two objects encountered at different time points. An intact temporal order memory is indicated by more time spent exploring the object encountered earlier compared to the newer one. The test was conducted in an arena measuring 43 x 43 cm. The animals were habituated in the same arena the day before for a duration of one hour. The task consisted of three sessions, each lasting five minutes, with one-hour intervals between the first two sessions and a three-hour interval between the second and the third sessions, as described in ^84^.

In the first and second sessions, two different couples of identical objects were introduced into the arena, and the animals were free to explore them. In the last session, one copy of the object from the first phase and one copy of the object from the second phase were placed in the arena. The position of objects in this phase was randomly selected and counterbalanced across the different groups. A discrimination index was computed as the disparity between the time spent exploring the object from the first phase compared to the object from the second phase, divided by the total time spent exploring both objects. A discrimination index exceeding 0 suggests intact temporal order memory. Subjects who did not complete at least 2 seconds of exploration during the experimental phases were excluded from the analysis. One animal was excluded for this reason. Another animal was exempted from testing due to health issues (corneal opacity). Data were analyzed using a 2-way ANOVA with genotype and treatment factors followed by a Sidak’s multiple comparisons test.

#### Partial Least Square Analyss

Partial least square correlation analyses (PLSC) ^44^ was used to assess the relationship between resting state functional connectivity and behavior in our mice cohort. PLSC is a multivariate method which finds brain-behavioral latent components (LC) that maximally co-vary while being uncorrelated to other LVs. The null hypothesis that the observed brain-behavior relationship could be due to chance was tested via permutation testing (1,000 iterations) of the behavioral data. To prevent LCs from being driven by inter-group differences, permutation was performed within groups ^85^. The stability of the contribution of each brain and behavioral element was assessed via bootstrapping with replacement (1,000 iterations), and also computed within experimental group to avoid covariance patterns being driven by group differences ^85^. PLS correlation was carried out between the behavior at each age and connectivity in default mode and ventral hippocampal network. These networks encompass brain regions implicated in the behaviors assessed ^10, 46,47^. Treatment was entered as a binary design variable in the behavioral matrix to assess the genotype-specific contribution of treatment to the observed brain-behavioral covariance.

### Human studies

#### Participants

The human sample used in this study comprised 256 participants, recruited from three research sites. The sample was divided into two cross-sectional cohorts: the first cohort consisted of individuals aged 6-11 years old, including 21 molecularly confirmed 22q11.2 deletion carriers (22q11DS) and 31 demographically comparable Healthy Controls (HC). The second cohort encompassed participants aged from puberty to 30 years (referred to hereafter as Post-puberty), comprising HC (n = 86) and 22q11DS (n = 118). As we lacked direct measures of pubertal onset, 11 years of age was chosen as a cut-off between childhood and Post-puberty groups across sexes on the basis of epidemiological studies estimating 10-11 as crucial ages for puberty onset ^86^. Deletion carriers and healthy controls were matched based on age and sex. Aggregate and site-specific demographics are reported in Table 1 and Supplemental Table S1, respectively.

Participants with neurological or medical conditions unrelated to 22q11.2 deletion, insufficient fluency in English, substance or alcohol abuse and/or dependence within the past 6 months, as well as HC individuals meeting diagnostic criteria for any major mental disorder upon administration of the Structured Clinical Interview for DSM-IV Axis I Disorder, were excluded from the study. Adult participants provided written consent, while participants under the age of 18 years required written consent from their parents or guardians. All procedures and informed consent documents received approval from the Institutional Review Board at each academic center.

#### Clinical assessment

The participants’ parents were assessed using the Social Responsiveness Scale (SRS) for children (6-17 years old) or adults (18 years and older) ^87^, in addition to an Autistic Diagnostic Interview (ADI). Verbal IQ and nonverbal IQ were evaluated using the Weschler Abbreviated Scale of Intelligence-2 (WASI-2), which included the WASI-2 Vocabulary and WASI Matrix Reasoning tests. The scores from these tests were combined to derive a WASI full-scale IQ ^88^, as detailed in Table 1. For the assessment of neuropsychiatric diagnoses and symptomatology, participants were administered the Structured Clinical Interview for DSM-IV Axis I Disorders (SCID), with an additional developmental disorders module ^89^, the Structured Interview for Psychosis-Risk Syndromes (SIPS ^90^), the Autistic Diagnostic Observation Schedule (ADOS, ^91^), and the Brief Psychiatric Rating Scale (see ^92^for recruitment and clinical assessment details).

#### Resting state fMRI acquisition

fMRI timeseries were obtained from three different sites: UCLA, SUNY, and KCL. At UCLA, the images were acquired using two different scanners. One subset of subjects from the UCLA sample (Childhood: HC = 21, 22q11DS n= 11; Post-puberty: HC = 24; 22q11DS n = 32) was scanned on a 3 Tesla Siemens Tim Trio Scanner equipped with a 12-channel head coil receiver. The acquisition parameters for these rsfMRI BOLD images were as follows: voxel size 3.0 x 3.0 x 4.0 mm, TE=30 ms, TR=2000 ms, flip angle 90°, matrix size 64 × 64, FOV=192, 34 axial slices, and a slice thickness of 4.0 mm for 152 volumes. The other subset from UCLA (Childhood: HC = 5, 22q11DS n = 7; Post-puberty HC n = 7, 22q11DS n = 21) was imaged on a 3 Tesla Siemens Magnetom Prisma Fit scanner equipped with a 32-channel head coil receiver. The acquisition parameters for this subset were: voxel size 2.0 x 2.0 x 2.0 mm, TE=37 ms, TR=800 ms, flip angle 52°, matrix size 104 × 104, FOV=208, 72 axial slices, and a slice thickness of 2.0 mm for 420 volumes. Subjects from the SUNY sample (Post-puberty HC n = 28; 22q11DS n = 44) were imaged on a 3 Tesla Siemens Tim Trio Syngo MR B17 scanner with an 8-channel head coil receiver. The acquisition parameters for rsfMRI BOLD images at SUNY were: voxel size 4.0 x 4.0 x 4.0 mm, TE=30 ms, TR=2000 ms, flip angle 90°, matrix size 64 × 64, FOV=256, 34 axial slices, and a slice thickness of 4.0 mm for 152 volumes. At KCL (Childhood: HC n = 5, 22q11DS n = 3; Post-puberty: HC n =27, 22q11 DS n = 21), the images were acquired using a 3 Tesla GE Signa Hdx scanner, with the following acquisition parameters: voxel size 4.0 x 4.0 x 4.0 mm, TE=30 ms, TR=2000 ms, flip angle 90°, matrix size 64 × 64, FOV=256, 34 axial slices, and a slice thickness of 4.0 mm for 152 volumes.

#### Resting state fMRI preprocessing

Images were preprocessed using the preconfigured “preprocessing” pipeline (https://github.com/FCP-INDI/C-PAC/blob/master/CPAC/resources/configs/pipeline_config_preproc.yml) of the Configurable Pipeline for the Analysis of Connectomes (CPAC; https://fcp-indi.github.io/) ^93^. Briefly, functional scans were slice-timing corrected and motion corrected using rigid-body, 6 parameter transformations. Nuisance variables regression was then performed on the motion corrected data including the regression of a 24-parameter model of motion (including the 6 motion parameters, their square, their derivatives, and the square of their derivatives), 5 noise signals identified using CompCor obtain from White Matter, and the mean CSF signal. Scans were then filtered using a bandpass filter only retaining frequencies between 0.01 and 0.1 Hz and normalized to MNI 152 template with linear and non-linear registration. Smoothing was performed using a Gaussian kernel of 6mm FWHM. fMRI volumes in which the sum of mean frame-wise displacement across the six rigid body parameters exceeded 0.5 mm were scrubbed. Participants with over 40% of frames scrubbed were excluded from further analyses.

#### fMRI connectivity analyses

The objective of these analyses was to assess whether the functional dysconnectivity signature observed in the mouse model is translationally conserved in the human cohort. We conducted the same functional analyses in the human dataset for both age groups to maximize cross-species comparability. Specifically, we used global connectivity mapping to create an unbiased mapping of functional connectivity differences within a 25% gray-matter probability mask. To minimize the impact of interscanner variability on our results, we harmonized our global connectivity maps using Combat, a state-of-the-art tool based on an empirical Bayes formulation that minimizes between-site variability ^94^. Between-group differences between 22q11DS and HC was then analyzed while controlling for the effects of sex and number of retained volumes, and family-wise error (FWER) cluster-corrected for multiple comparisons as implemented in FSL (|*t*| > 2.0, p < 0.05), for each of the two age groups separately. Two-way ANOVA was then used to test the interaction between age group (Childhood, Post-puberty) and genotype (HC, 22q11DS). Similarly to the mouse model, we employed seeds placed in foci of altered global functional connectivity, as illustrated in Supplemental Figure S5, to compute seed-based functional connectivity. We then calculated between-group differences and the age x genotype interaction of these seed-based maps using the same method used for global functional connectivity. Non-linear developmental trajectories of global connectivity in areas undergoing significant reconfiguration were characterized using general additive mixed models (GAMMs, ^34^). A GAMM was fit to predict global connectivity from the smoothed effect of age and genotype for each area of interest (opercular cortex, postcentral gyrus and hippocampus), resulting in separate smooths for each group, 22q11DS and HC. Ages were global connectivity between the two genotypes significantly differed were defined as age ranges where the 95% confidence interval of the genotype difference in age smooths did not include zero.

For the subset of 22q11 deletion carriers with available Social Responsiveness Scale (SRS) scores (Childhood N = 18, Post-puberty N = 111), the relationship between 22q11DS-related functional dysconnectivity and autism-related behaviors was assessed in two ways. First, we probed this relationship for global connectivity of areas undergoing a significant reconfiguration over age (postcentral gyrus and opercular cortex) and run a linear model with average global connectivity in these ROIs as a predictor of SRS score for the two age groups separately (Childhood and Post-puberty). Secondly, we extended our analysis by using seed-based functional connectivity maps of foci of significant age x genotype interaction (postcentral gyrus and opercular cortex) to assess whether they would be more sensitive to behaviorally-relevant correlations. To assess which areas showed age-specific relationship between brain and behavior, we used a voxelwise linear model with seed-based functional connectivity of either seed as a predictor of SRS scores and assessed its interaction with age group, while controlling for the effects of sex and retained volumes. By including the interaction between functional connectivity and age group in the model, we were able to assess for which areas the relationship between connectivity and SRS followed an age-specific pattern.

#### Gene expression decoding and enrichment analysis

To probe which molecular mechanism might be involved in functional dysconnectivity of 22q11DS patients, we carried out a gene decoding and enrichment analysis. Gene expression decoding, as implemented by NeuroVault ^95^, aims at identifying genes whose spatial expression in the brain is similar to spatial patterns of functional connectivity. Briefly, a linear model is used to estimate spatial similarity between pattern of functional connectivity maps and individual gene expression maps for each of the 20,787 genes of the Allen Institute Human Brain Gene Expression atlas in each of the 6 donor’s brain. Gene and donor-specific slopes are then assessed using a one-sample t tests. The results of this t test are then multiple comparison corrected, and only the genes surviving FDR at *q* < 0.05 are used in the following analyses. In the current study, areas undergoing hyper-to-hypo reconfiguration over development (i.e., 22q11-related dysconnectivity) are represented by positive *t* values in our voxelwise global connectivity age x genotype interaction *t* maps. Hence, we restricted our search to genes which were positively correlated with our age x genotype interaction map (i.e., genes that were more expressed in areas of functional dysconnectivity). This resulted in a gene list of *N* = 3897 decoded genes, whose spatial expression positively correlates with areas of hyper-to-hypo reconfiguration in 22q11DS.

Firstly, to test the hypothesis that synaptic mechanisms could be implicated in the observed dysconnectivity in human 22q11DS too, we performed gene enrichment analyses between the decoded genes and GO Terms identifying genes implicated in synaptic signaling, as reported by ^49^. Furthermore, we also tested the hypothesis of a significant enrichment between decoded genes and GO Terms related to glial-related synaptic remodeling, given their involvement in this process ^39^.

Next, we set out to fully ascertain the parallelism between the dysconnectivity we observed in LgDel mice and 22q11 deletion carriers, specifically by looking at whether the decoded genes would be enriched for GSK3β-related synaptic genes. To do this, we obtained a broad list of GSK3β interacting genes using protein-protein interaction analysis in STRING-DB^96^, with a medium confidence setting (0.4) and allowing up to 500 interactors. Given the broadness of this initial list and mouse-derived evidence of a mechanistic involvement of synaptic processes in functional dysconnectivity, we then increased its specificity by pruning the initial GSK3β-interactors to only include genes involved in synaptic processes, as per our hypothesis guided by dendritic spine investigations. Specifically, we used the Syngo ^50^ synaptic gene list (which includes genes involved in biological processes in the presynapse and in the post-synapse, synaptic metabolism, synaptic signaling, transport, synaptic organization) to reduce the initial *n* = 501 GSK3β-interactors to *n* = 113 GSK3β-synaptic interactors. Furthermore, we used a generative null model to assess the specificity of the observed enrichment for the spatial topography of our developmental dysconnectivity pattern. To this aim, we used a spatial autocorrelation preserving generative null model ^97^ to generate k=1000 surrogate maps whose spatial autocorrelation matched that of the empirical developmental dysconnectivity (i.e., global connectivity age x genotype interaction) map. We then applied gene decoding to each surrogate map, and performed gene enrichment analysis between the resulting decoded genes and the synaptic interactors of GSK3B, yielding 1000 overlap sizes. We then compared the number of genes overlapping for our empirical developmental dysconnectivity map with the generated null distribution. Finally, we exploited gene enrichment analyses to assess the clinical relevance of the functional dysconnectivity signature we observed. Specifically, we performed gene enrichment analyses between the decoded genes and previously published disorder-relevant gene lists (^51, 52, 98, 99^ and https://gene.sfari.org, downloaded in April 2023).

Prior to all enrichment analyses, all gene lists were pruned to only include protein coding genes expressed in the brain (*N* = 16796 genes, ^51^), which was hence considered as the background pool. For all gene enrichment analyses, odds ratio and hypergeometric p values for enrichment analyses were computed using code publicly available at https://github.com/mvlombardo/utils/blob/master/genelistOverlap.R. FDR correction was applied to correct for multiple comparison where appropriate, and corrected *q* values are reported for all enrichments.

---

**Supplementary Table 1.** Site-specific subject demographics.

|  | Children<br>(6-11 years) |  |  | Post-pubertal<br>(12-30 years) |  |  |
| --- | --- | --- | --- | --- | --- | --- |
| UCLA (Trio) | HC (n=21) | 22q11DS<br>(n=11) | *p value* | HC (n=24) | 22q11DS (n=32) | *p value* |
| Age (years) | 8.6 ± 1.9 | 9.3 ± 2 | 0.34 | 16.6±3.6 | 17.6±3.9 | 0.31 |
| Sex (M/F) | 11/10 | 6/5 | >0.99 | 12/12 | 12/20 | 0.4 |
| Full scale IQ* (mean ± SD) | 117.1±<br>26.1 | 78.2± 8.8 | <0.0001 | 106± 16.6 | 79.9± 14.7 | <0.0001 |
| ASD, n (%) | 0 (0) | 7 (64) | <0.0001 | 0 (0) | 14 (44) | <0.0001 |
| ADHD, n (%) | 1 (5) | 8 (73) | <0.0001 | 1 (4) | 14 (44) | 0.0008 |
| Psychotic disorder, n (%) | 0(0) | 0 (0) | - | 0 (0) | 3 (9.4) | 0.25 |
| Anxiety disorder, n (%) | 1(5) | 4 (36) | 0.04 | 7 (29) | 18 (56) | 0.06 |
| Mood disorder, n (%) | 0 (0) | 1 (9) | 0.34 | 4 (17) | 8 (25) | 0.5 |
| Current antipsychotic<br>treatment, n (%) | 0 (0) | 0 (0) | - | 0 (0) | 5 (16) | 0.06 |
| Current<br>antidepressant/antianxiety<br>treatment, n (%) | 0 (0) | 0 (0) | - | 0 (0) | 9 (28) | 0.007 |
| Current psychostimulant<br>treatment, n (%) | 1 (5) | 1 (9) | 0.34 | 0 (0) | 7 (22) | 0.02 |
| UCLA (Prisma) | HC (n=5) | 22q11DS (n=7) | *p value* | HC<br>(n=7) | 22q11DS (n=21) | *p value* |
| Age (years) | 9.9 ± 1.9 | 9.6 ± 1.7 | 0.8 | 15.3 ±<br>2.83 | 19.4 ± 4.4 | 0.04 |
| Sex (M/F) | 1/4 | 2/5 | >0.99 | 2/5 | 6/15 | >0.99 |
| Full scale IQ* (mean ± SD) | 120.4±<br>14.4 | 76.1± 5.7 | <0.0001 | 118 ±<br>20.5 | 86.6 ± 12.8 | <0.0001 |

| ASD, n (%) | 0 (0) | 2 (29) | 0.47 | 0 (0) | 6 (29) | 0.29 |
| --- | --- | --- | --- | --- | --- | --- |
| ADHD, n (%) | 0 (0) | 1 (14) | >0.99 | 1 (14) | 9 (43) | 0.36 |
| Psychotic disorder, n (%) | 0 (0) | 0 (0) | - | 0 (0) | 1 (4.8) | >0.99 |
| Anxiety disorder, n (%) | 0 (0) | 2 (29) | 0.47 | 0 (0) | 9 (43) | 0.06 |
| Mood disorder, n (%) | 0 (0) | 1 (14) | >0.99 | 0 (0) | 4 (19) | 0.54 |
| Current antipsychotic treatment, n (%) | 0 (0) | 1 (14) | >0.99 | 0 (0) | 1 (4.8) | >0.99 |
| Current antidepressant/antianxiety treatment, n (%) | 0 (0) | 0 (0) | - | 0 (0) | 2 (9.5) | >0.99 |
| Current psychostimulant treatment, n (%) | 0 (0) | 0 (0) | - | 1 (14) | 3 (14) | >0.99 |
| KCL | HC (n=5) | 22q11DS (n=3) | *p value* | HC (n=27) | 22q11DS (n=21) | *p value* |
| Age (years) | 8.8±1.3 | 9.3±2.1 | 0.66 | 18.5±5.4 | 18.3±4.6 | 0.9 |
| Sex (M/F) | 3/2 | 1/2 | >0.99 | 12/15 | 10/11 | >0.99 |
| Full scale IQ* (mean ± SD) | 114.8 ± 10 | 80.5 ± 2 | 0.01 | 111.1 ± 14.9 | 83.4 ± 14.4 | < 0.0001 |
| ASD, n (%) | 0 (0) | 0 (0) | - | 0 (0) | 0 (0) | - |
| ADHD, n (%) | 0 (0) | 1 (33) | 0.37 | 0 (0) | 2 (9.5) | 0.2 |
| Psychotic disorder, n (%) | 0 (0) | 0 (0) | - | 0 (0) | 1 (4.8) | 0.44 |
| Anxiety disorder, n (%) | 0 (0) | 1(33) | 0.37 | 0 (0) | 10 (47.6) | < 0.0001 |
| Mood disorder, n (%) | 0 (0) | 0 (0) | - | 0 (0) | 1 (4.8) | 0.44 |
| Current antipsychotic treatment, n (%) | 0 (0) | 0 (0) | - | 0 (0) | 3 (14.3) | 0.08 |
| Current antidepressant/antianxiety treatment, n (%) | 0 (0) | 0 (0) | - | 0 (0) | 2 (9.5) | 0.2 |
| Current psychostimulant treatment, n (%) | 0 (0) | 0 (0) | - | 0 (0) | 0 (0) | - |
| **SUNY** | **HC (n=0)** | **22q11DS (n=0)** | **-** | **HC (n=28)** | **22q11DS (n=44)** | ***p value*** |
| Age (years) | - | - | - | 20.8± 1.4 | 21.1±2.3 | 0.5 |
| Sex (M/F) | - | - | - | 15/13 | 26/18 | 0.8 |
| Full scale IQ* (mean ± SD) | - | - | - | 108.5±16 | 75.1±12.1 | <0.0001 |
| ASD, n (%) | - | - | - | 0 (0) | 0 (0) | - |
| ADHD, n (%) | - | - | - | 5 (17.8) | 15 (34) | 0.2 |
| Psychotic disorder, n (%) | - | - | - | 0 (0) | 2 (4.5) | 0.52 |
| Anxiety disorder, n (%) | - | - | - | 3 (10.7) | 12 (27.3) | 0.1 |
| Mood disorder, n (%) | - | - | - | 1 (3.6) | 13 (29.5) | 0.0063 |
| Current antipsychotic treatment, n (%) | - | - | - | 0 (0) | 3 (6.8) | 0.3 |
| Current antidepressant/antianxiety treatment, n (%) | - | - | - | 2 (7) | 11 (25) | 0.07 |
| Current psychostimulant treatment, n (%) | - | - | - | 2 (7) | 6 (14) | 0.5 |
| * IQ data were not available for 1 child 22q11DS, 2 Post-pubertal HC in TRIO; 1 children HC, 1 children 22q11DS, 6 Post-pubertal HC, 2 Post-pubertal 22q11DS in KCL, 1 Post-pubertal HC. |  |  |  |  |  |  |

