## Supplementary Figures for "Synaptic-dependent developmental dysconnectivity in 22q11.2 deletion syndrome"

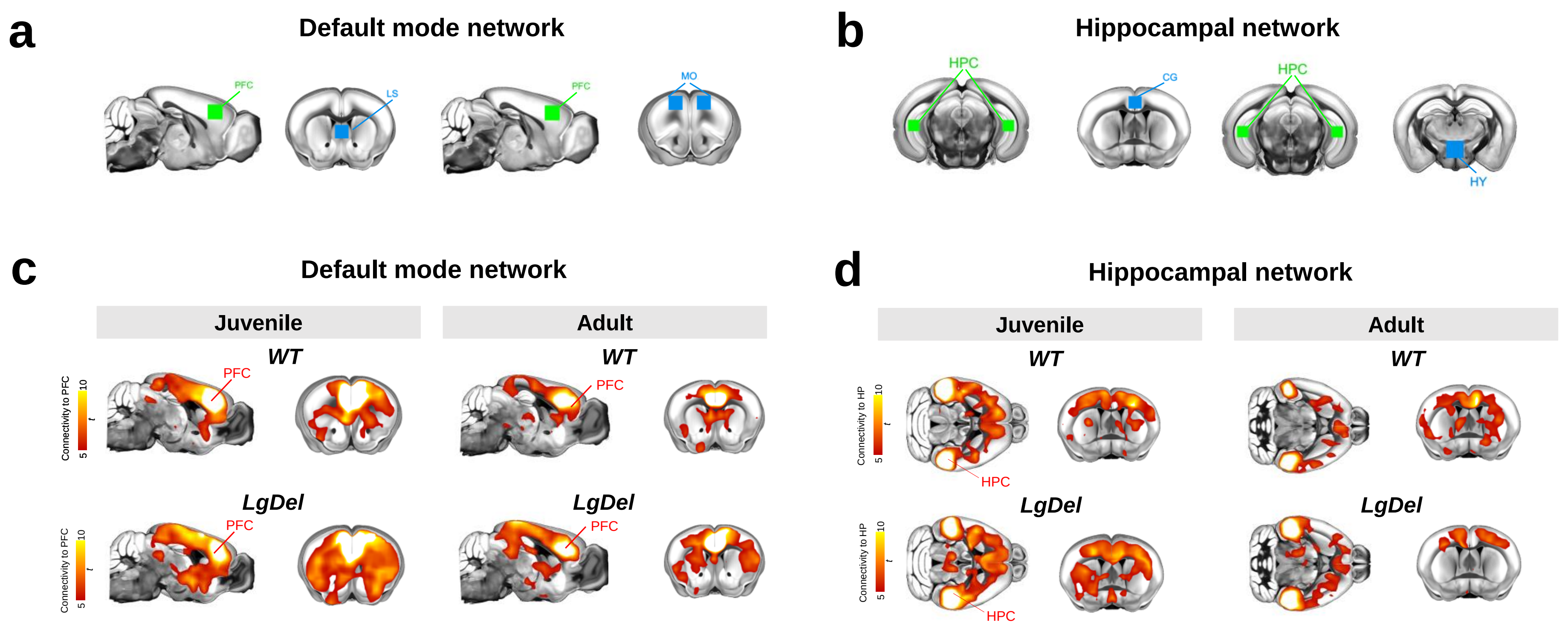


Figure S1. Seed-based mapping of DMN and hippocampal networks in juvenile and adult (post pubertal) LgDel mice. *(a, b) Location of seed (in green) used for seed-based probing of the DMN and hippocampal networks. Network connectivity was carried out by computing pairwise connectivity between the seed and the ROIs indicated in blue (b, c) Seed-based fMRI connectivity maps of the DMN and hippocampal networks in WT and LgDel mice at juvenile and adult stage. [CG, Cingulate cortex, LS, lateral septum, MO, motor cortex, HY, hypothalamus, PFC, prefrontal cortex, HPC, hippocampus].*

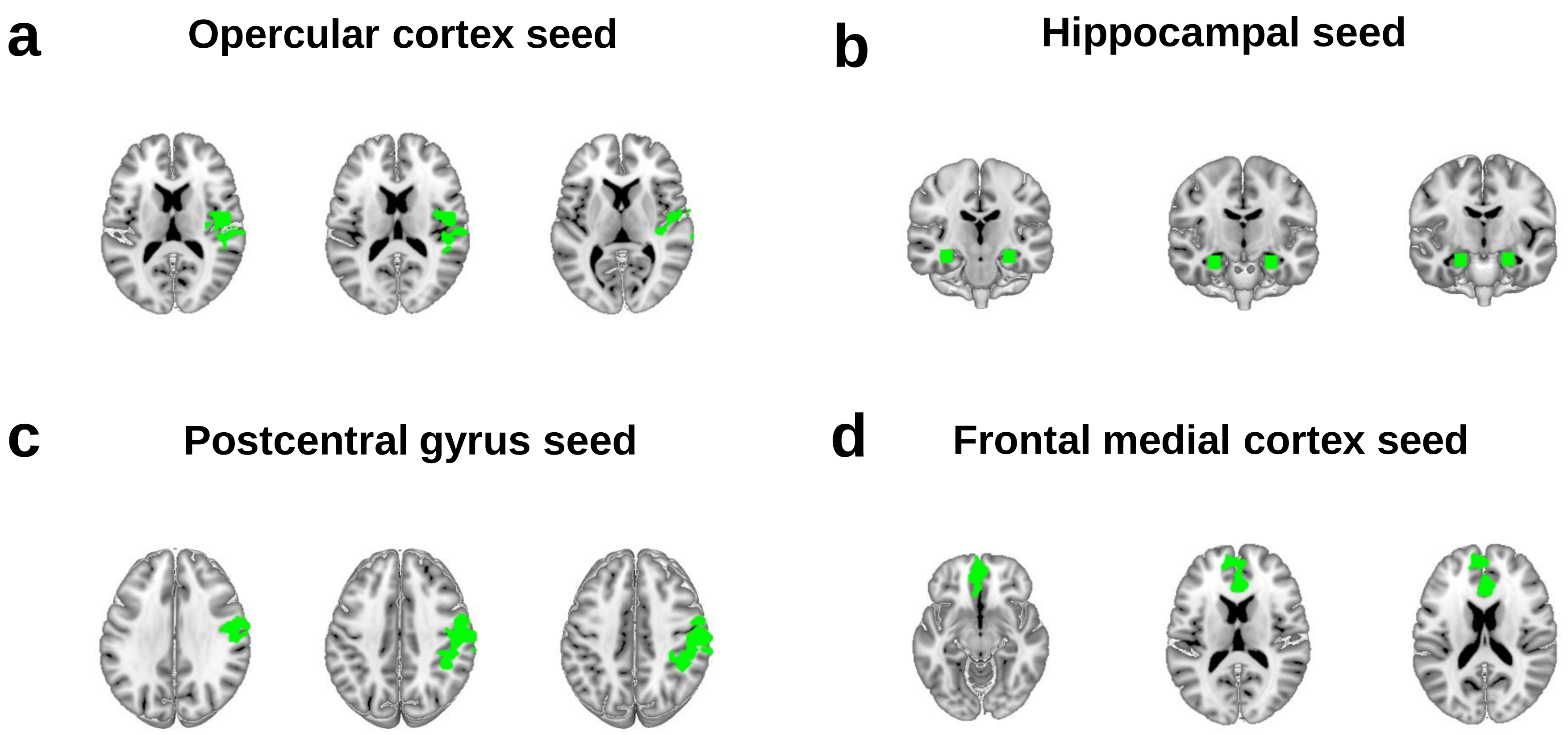


Figure S2. *Location of seed used to probe fMRI connectivity in our investigations of human 22q11DS: opercular cortex (a), hippocampus (b), postcentral gyrus (c) and frontal medial cortex (d). Seeds in a and b reconstitute extension of significant genotype x age interaction clusters in fMRI global connectivity analysis. Hippocampal seed was manually designed as a cubic region of interest (ROI) (3 x 3 x 3 voxels) bilaterally centered over the hippocampus. Frontal medial cortex seed reconstitutes extension of prefrontal hypoconnectivity cluster in fMRI global connectivity analyses of adult group.*

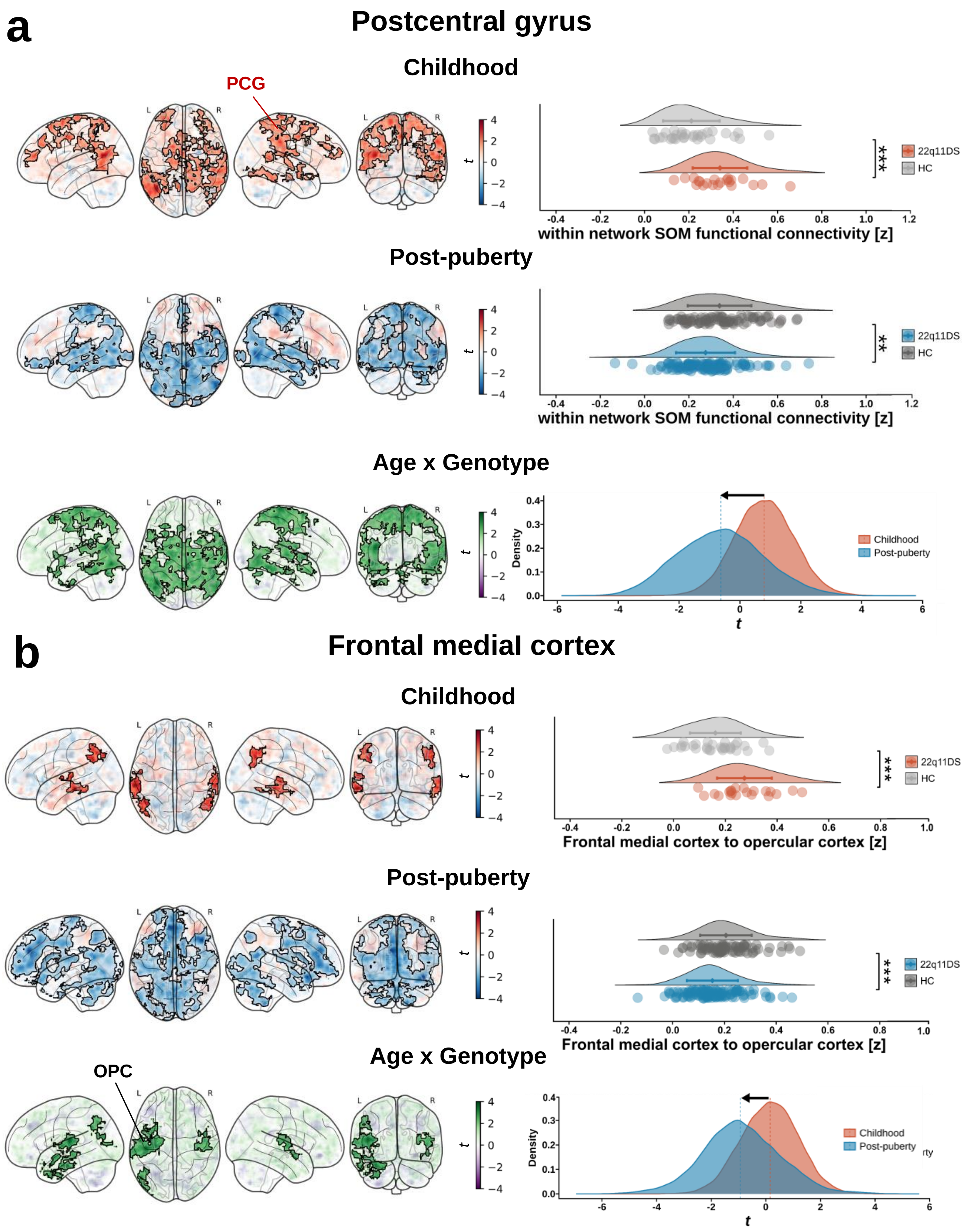


Figure S3. Seed-based analysis of functional connectivity of postcentral gyrus and frontal medial cortex. *(a) Voxel-wise (left panels) mapping of seed-based fMRI connectivity of the postcentral gyrus (a) and frontal medial cortex (b) revealed increased functional connectivity in 22q11DS individuals during childhood; in contrast, 22q11DS showed reduced fMRI connectivity relative to controls in the post-pubertal cohort. Semi-transparent maps in the background illustrate unthresholded t-values. Clusters surviving FWER correction (|t|>2.0, p < 0.05) are outlined in black. Areas exhibiting a significant age x genotype interactions were identified using a linear model. Whole-brain distribution of t values resulting from group differences at each age revealed a robust shift (arrow) from prevalent hyperconnectivity in childhood to prevalent hypoconnectivity after puberty (bottom panel). [PCG, postcentral gyrus, OPC, opercular cortex].*

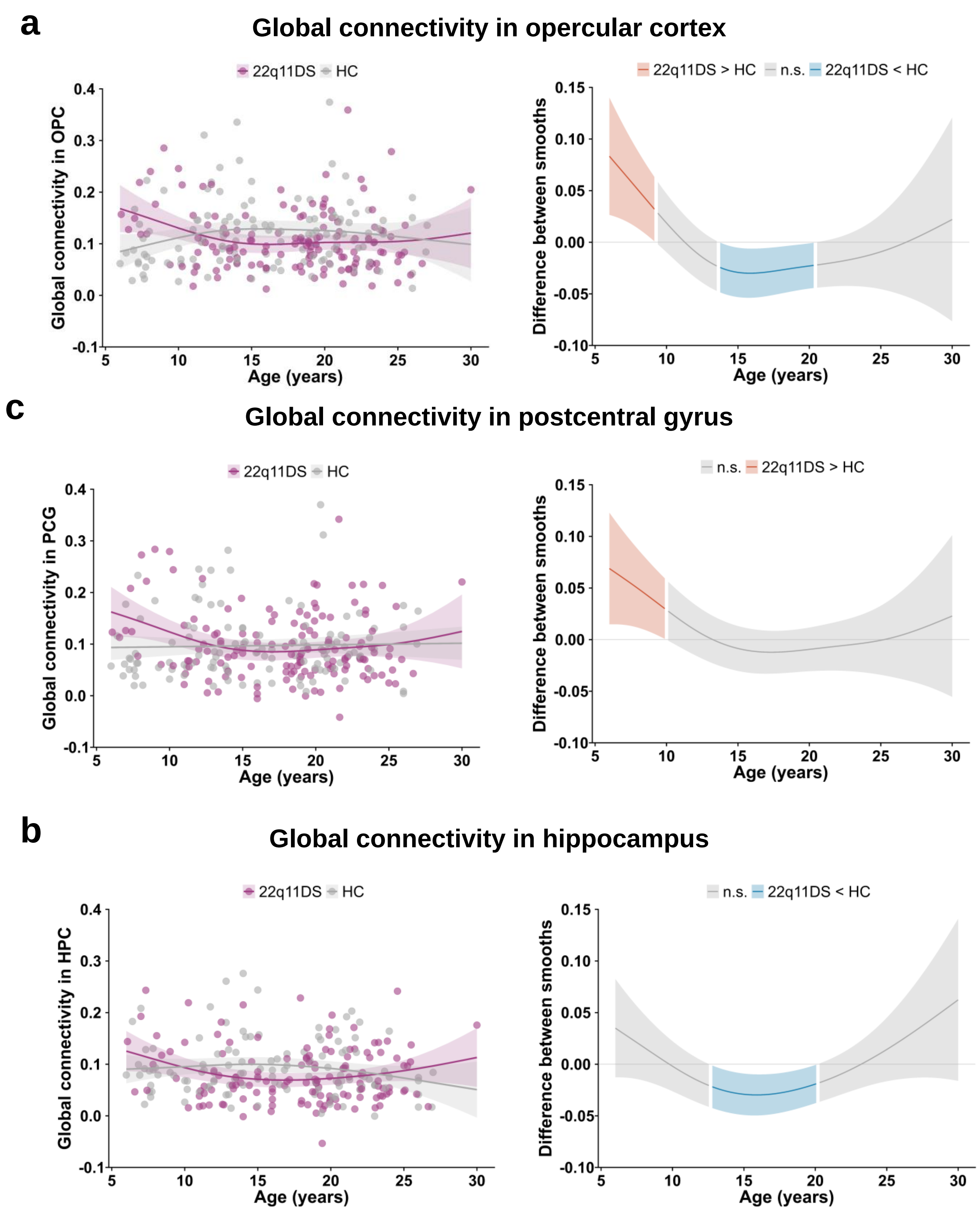


Figure S4. Developmental trajectories of global fMRI connectivity in opercular cortex, postcentral gyrus and hippocampus as probed with a non-linear mixed model

*Smoothed age curves (left) for global connectivity in opercular cortex (a), postcentral gyrus (b) and hippocampus (c) from generalized additive mixed effect models (GAMMs). The model predicts connectivity from age and genotype. Intergroup comparisons (right) computed as difference between smooths show that global hyperconnectivity relative to controls characterizes 22q11 deletion carriers until late childhood (a, b), before reverting to hypoconnectivity during adolescence (a, c). [HPC, hippocampus, PCG, postcentral gyrus, OPC, opercular cortex].*

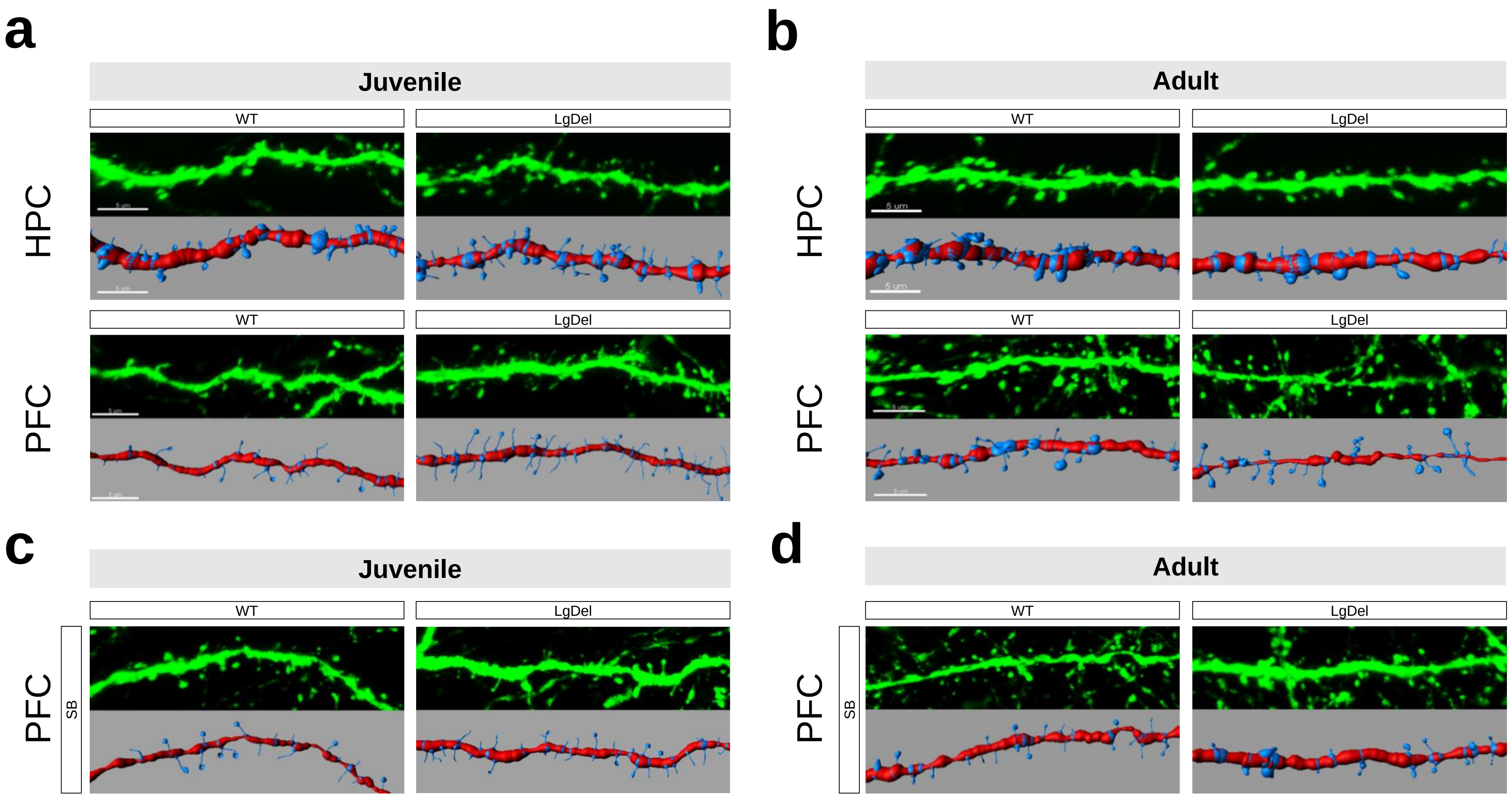


Figure S5. Alterations in dendritic spine density in LgDel mice and their rescue by SB. *Panel illustrating representative GFP-labeled neurons and corresponding 3D reconstruction of dendritic spines in hippocampus (HPC) and prefrontal cortex (PFC) of juvenile (a) and adult (b) WT and LgDel vehicle-treated mice. Panels (c) and (d) show representative PFC neurons from SB-treated juvenile (c) and adult (d) WT and LgDel mice.*

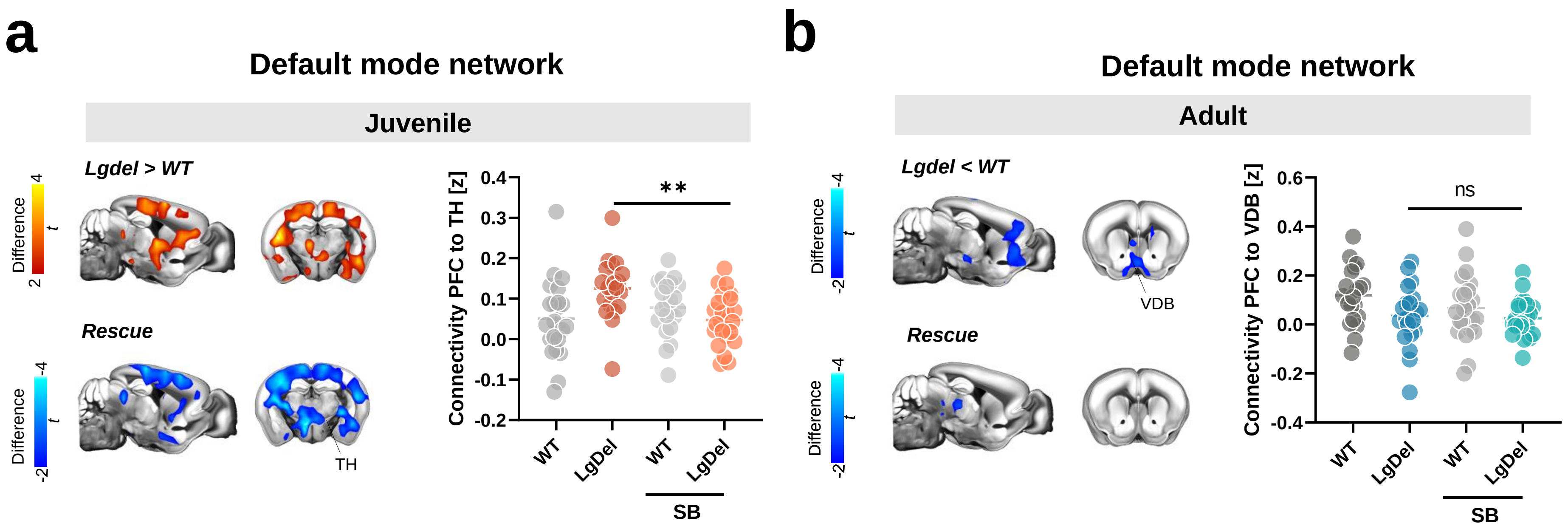


Figure S6. Seed-based probing of DMN connectivity in LgDel mice receiving SB treatment. *(a) Seed-based probing of DMN connectivity in juvenile mice revealed normalization of fMRI hyperconnectivity by developmental SB-treatment. (b) The same treatment did not ameliorate hypoconnectivity of basal forebrain areas in adult LgDel mice [PFC, prefrontal cortex, TH, thalamus, VDB, ventral diagonal band].*

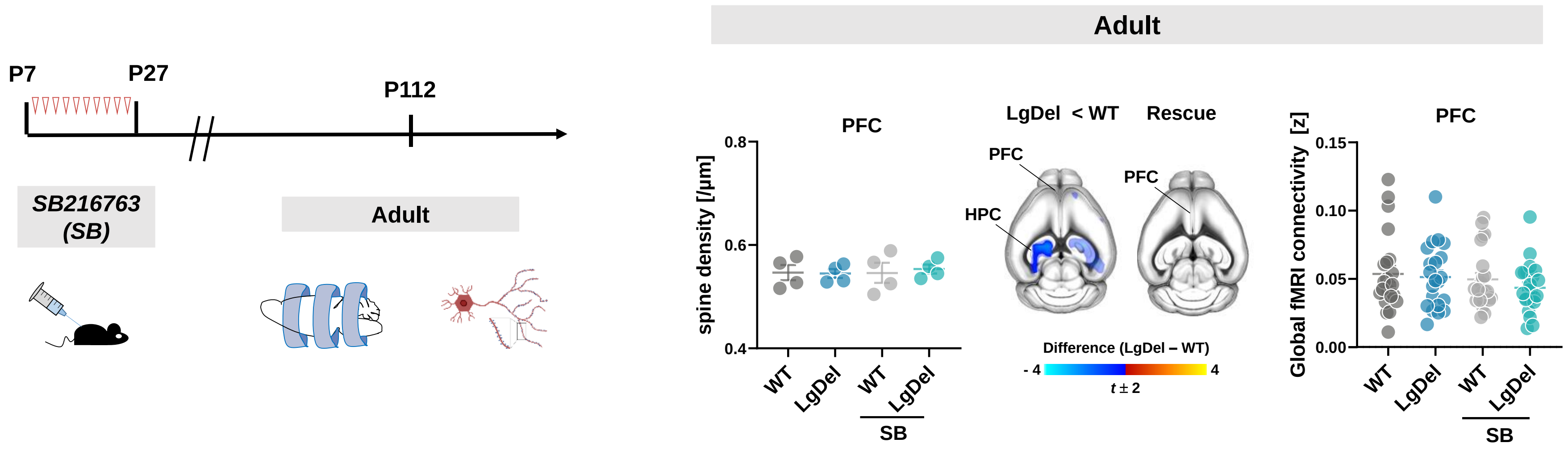


Figure S7. Synaptic counts and fMRI dysconnectivity in prefrontal cortex (PFC) of adult LgDel mice. *Left: Experimental timeline of the GSK3β inhibition treatment protocol and imaging and spine count studies. Right: Concordant lack of synaptic and connectivity alterations in PFC of adult mice. SB-treatment did not affect dendritic spine density at this age [HPC, hippocampus, PFC, prefrontal cortex].*

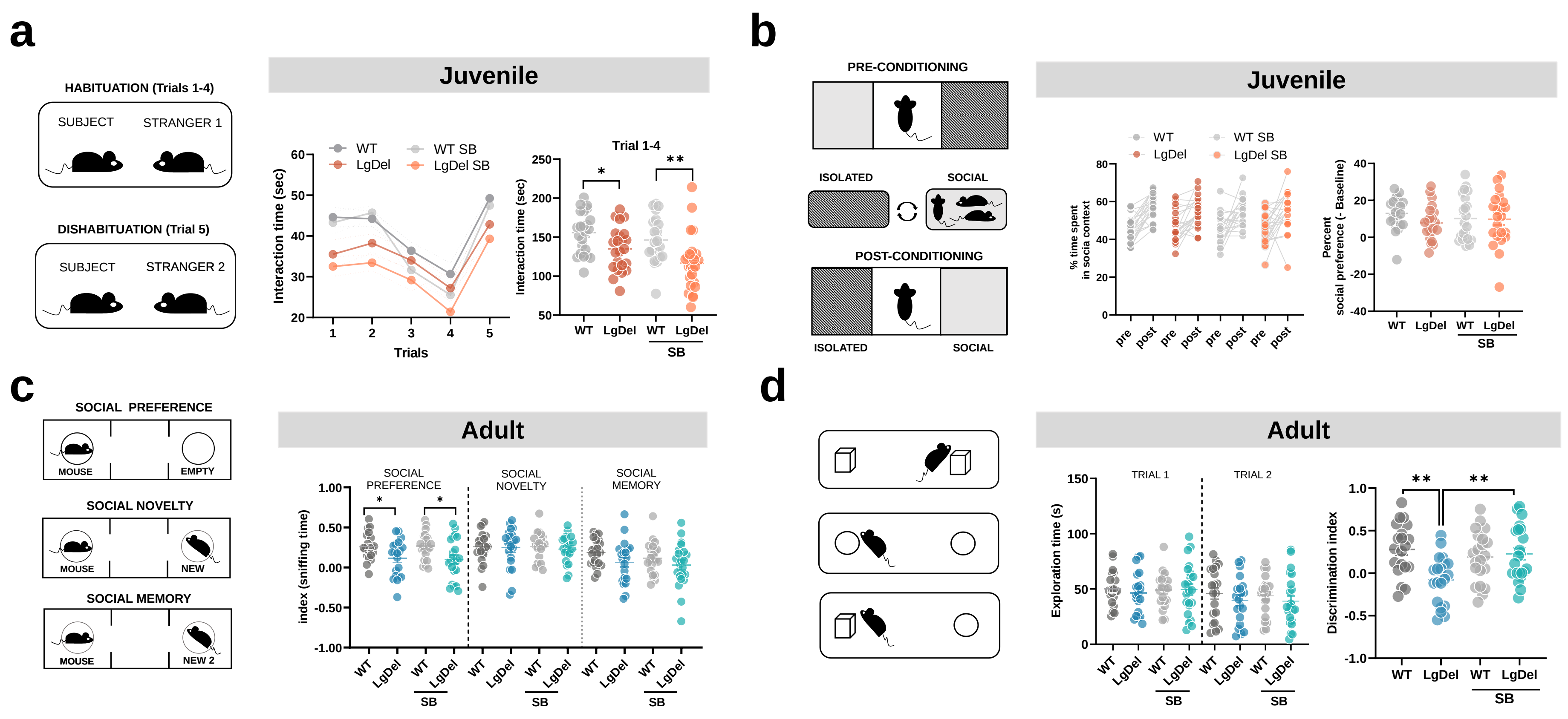


Figure S8. Behavioral assessments in juvenile and adult LgDel mice. a) *Scheme of the habituation-dishabituation social interaction task (left). Reduced interaction time in juvenile LgDel mice was not rescued by pharmacological GSK3β inhibition (SB). In LgDel mice, reduced interaction time was observed across trials, thereby suggesting an absence of social recognition deficits.* b) *Scheme of the social conditioning place preference task (SCPP) (left). LgDel mice did not display impairments in social reward learning. c) Scheme of the three-chambered social approach task (left). Comparison between groups measured as difference in sniffing time of cups in two compartments (index). LgDel mice displayed a reduced sociability index in the social preference phase which was not rescued by SB treatment. No intergroup differences were detected in the social novelty or social memory phase either. d) Scheme of the temporal order memory task (left). A robust impairment in temporal order memory was observed in adult LgDel mice. This impairment was robustly rescued by developmental administration of SB (plot on the right). *$p<0.05$, **$p < 0.01$.*

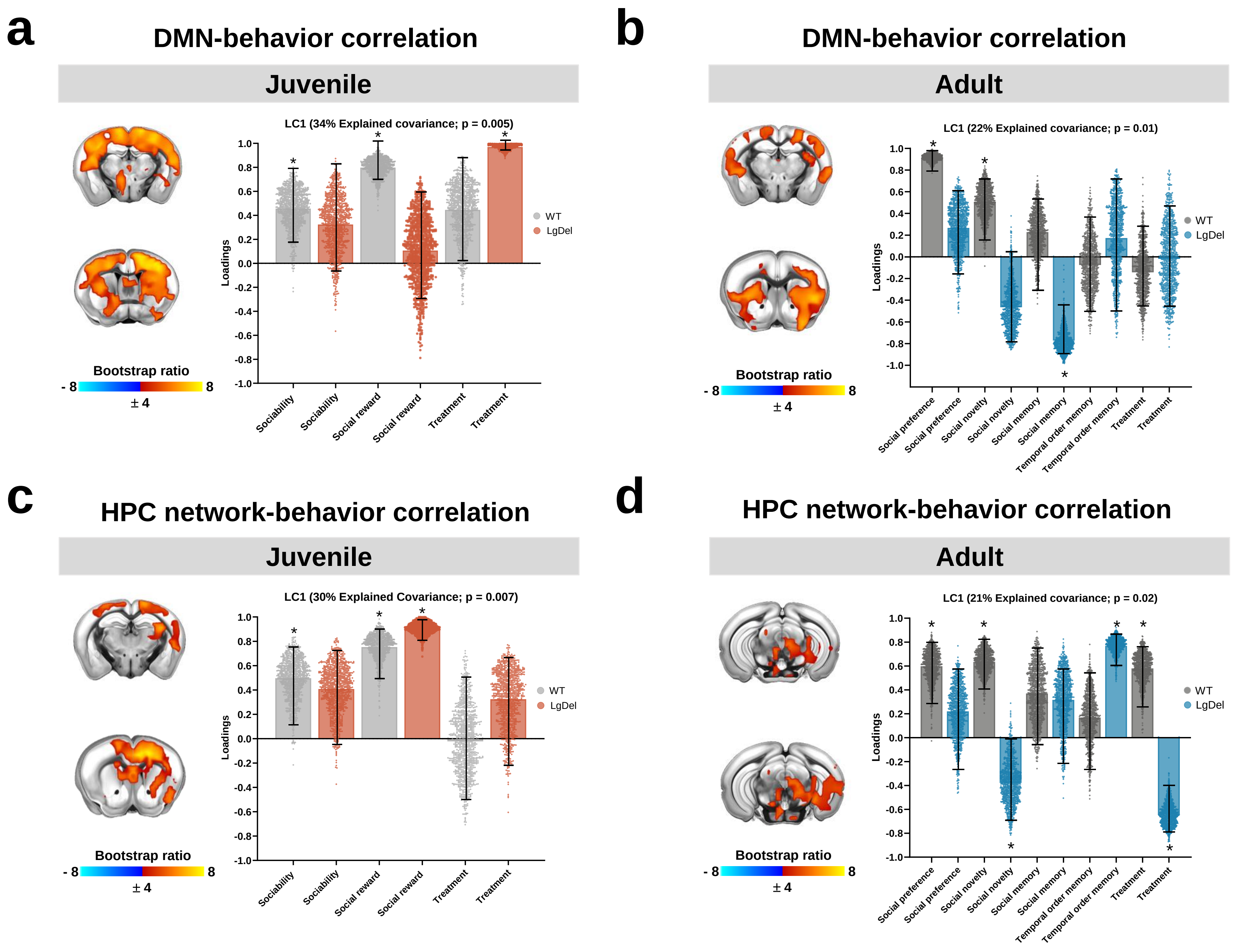


Figure S9. Disrupted relationship between resting state fMRI connectivity and behavior in LgDel mice. *Partial least squares (PLS) correlation analysis identified latent components capturing the relationship between prefrontal (a, b) or hippocampal (c, d) connectivity and multivariate socio-behavioral patterns, in both juvenile and adult mice. This analysis revealed that PFC connectivity to sensory and striatal regions was associated with sociability and social reward in wildtype mice across ages (a, b). This relationship was disrupted in LgDel mice, in which only GSK3β inhibition (a, encoded categorically) or social memory (b) were associated with the observed patterns of prefrontal connectivity. (c) Hippocampal connectivity to prefrontal or sensory regions was associated with sociability and social reward in juvenile wildtype mice. This relationship was only partially preserved in juvenile LgDel mice. (d) Hippocampal connectivity to midbrain was associated with sociability as well as GSK3β treatment in wildtype mice. In LgDel mice, this relationship was disrupted. Specifically, the association between connectivity pattern and social novelty or treatment in LgDel appeared to be reversed in terms of directionality as compared to wildtype group. Hippocampal connectivity to midbrain was also associated with temporal order memory in the LgDel group only. Errors bars indicate 95% Confidence interval estimated by bootstrapping. Asterisks indicate stability of the variable (95% confidence intervals not including zero) [HPC, hippocampus].*
